# Rabaptin5 targets autophagy to damaged endosomes and SCVs by interaction with FIP200 and ATG16L1

**DOI:** 10.1101/2020.09.01.277764

**Authors:** Valentina Millarte, Simon Schlienger, Simone Kälin, Martin Spiess

## Abstract

Selective autophagy of damaged organelles is an important process for cellular homeostasis. The mechanisms how autophagy selects specific targets is often poorly understood. Rabaptin5 was previously known as a major regulator of early endosome identity and maturation. Here we identified two novel Rabaptin5 interactors: FIP200, a subunit of the ULK1 autophagy initiator complex, and ATG16L1, a central component of the E3-like enzyme in LC3 lipidation. Indeed, autophagy of early endosomes damaged by chloroquine or monensin treatment was found to require Rabaptin5 and particularly a specific short sequence motif binding to the WD domain of ATG16L1. Rabaptin5 and this interaction with ATG16L1 is further required for much of autophagic elimination of *Salmonella enterica* in phagosomes with early endosomal characteristics early after infection. Our results demonstrate a novel function of Rabaptin5 in quality control of early endosomes in the selective recruitment of autophagy to damaged early endosomes and phagosomes.

## INTRODUCTION

Endosomes are dynamic organelles that receive endocytic cargo from the plasma membrane and exocytic material from the trans-Golgi for sorting to late endosomes and lysosomes, to the cell surface via recycling endosomes, or back to the Golgi (Naslavsky and Caplan, 2018). Endosomal identities are defined by specific Rab GTPases, their effectors, and characteristic phosphoinositides. At early endosomes, Rab5 is the hallmark GTPase that activates VPS34/p150 to produce phosphatidylinositol-3-phosphate (PI3P) and recruits early endosome antigen 1 (EEA1) and Rabenosyn-5, two multivalent PI3P-binding proteins that act as membrane tethers to mediate homotypic endosome fusion. Rab5·GTP and PI3P are also responsible for recruitment of the Mon1/Ccz1 complex to activate Rab7 and deactivate Rab5 in the process of Rab conversion during maturation from early to late endosomes (Huotari and Helenius, 2011; Poteryaev et al., 2010).

Rab5 activity is regulated by a complex of Rabaptin5 and Rabex5, the GDP/GTP exchange factor of Rab5. Rabaptin5 binds to Rab4·GTP and Rab5·GTP, and Rabex5 binds to ubiquitin to mediate membrane recruitment and to activate Rab5 on early endosomes (Kälin et al., 2016; 2015; Mattera and Bonifacino, 2008; Mattera et al., 2006).

Searching for new interactors of Rabaptin5, we performed a yeast two-hybrid screen and discovered as a novel binding partner FIP200, a component of the ULK1–FIP200–ATG13–ATG101 autophagy initiator complex. Autophagy is a self-degradative survival mechanism of eukaryotic cells important to preserve cellular homeostasis in response to stress conditions such as lack of nutrients, accumulation of misfolded or aggregated proteins, damaged organelles, and pathogen infection (Bento et al., 2016; Dikic and Elazar, 2018; Mercer et al., 2018; Morishita and Mizushima, 2019). Clearance of damaged organelles is an important process to preserve overall organelle function. Defects in its mechanisms have been shown to contribute to some forms of cancer, to neurodegeneration, and inflammatory diseases (Anding and Baehrecke, 2017; Leidal et al., 2018). Selective autophagy has been described for many organelles including mitochondria, peroxisomes, endoplasmic reticulum, and lysosomes (mitophagy, pexophagy, ER-phagy, and lysophagy; Kirkin and Rogov, 2019).

Typically, autophagy is set in motion by activation of the ULK1–FIP200 complex under the control of mTOR. The ULK complex initiates a dynamic interactome of autophagy components by recruiting and activating the class III PI 3-kinase complex VPS34–VPS15–Beclin1–ATG14 to produce PI3P, further recruiting PI3P-binding proteins, such as WIPI (WD-repeat phosphoinositide interacting protein) family members. Together, these complexes and PI3P bind ATG16L1 with its partners ATG12–ATG5 acting as an E3-ubiquitin ligase-like enzyme for the lipidation of LC3 family proteins for elongation and closure of the phagophore membrane and fusion with lysosomes.

Selective autophagy of organelles destined for degradation differs from starvation-induced autophagy in its mechanism of initiation, which is independent of the nutrient situation, and in its targeting to specific cargo (Kirkin and Rogov, 2019). Autophagy is often recruited to damaged organelles after they are marked for degradation by ubiquitination of lumenal proteins or by galectins binding to exposed glycans (Randow and Youle, 2014). Galectins 3, 8, and 9 have been implicated in the recognition of damaged lysosomes, endosomes, or salmonella-containing vacuoles together with galectin-interacting autophagy receptors NDP52 or TRIM16 (Chauhan et al., 2016; Fraser et al., 2019; Jia et al., 2018; 2020; Thurston et al., 2012).

ATG16L1 emerges as an important hub for specific cargo to connect to the autophagy machinery (reviewed by Gammoh, 2020). Besides its role in LC3 lipidation, it has binding sites for ubiquitin, FIP200 and WIPI2, phosphoinositides, and generally for membranes. It was further shown to interact with TRIM16 on damaged lysosomes (Chauhan et al., 2016), and with TMEM59 involved in autophagic degradation of late endosomes/lysosomes and *Staphylococcus aureus*-containing phagosomes (Boada-Romero et al., 2013).

ATG16L1 is also involved in LC3-associated phagocytosis (LAP), where the LC3 conjugation system is recruited directly to a single-membrane phagosome for fusion with lysosomes (Heckmann et al., 2017; Martinez et al., 2011; 2015). LAP has been described mainly in phagocytes like macrophages, microglia, and dendritic cells that actively take up bacteria, opsonized particles, or dead cells by specific cargo receptors (such as toll-like receptors, immunoglobulin receptors, and the phosphatidylserine receptor TIM4, respectively). LAP is independent of the ULK–FIP200–ATG13–ATG101 complex and of WIPI proteins.

So far, little is known about selective autophagy of damaged early endosomes. Autophagy of endosomes disrupted upon uptake of transfection reagent-coated latex beads showed a requirement for ubiquitination and ATG16L1-binding to ubiquitin (Fujita et al., 2013). As an alternative method of damaging endosomes, lysosomotropic agents like chloroquine and monensin have been used, which cause swelling of acidic compartments due to osmotic imbalance (Florey et al., 2015; Jacquin et al., 2017; Mauthe et al., 2018). They induce autophagy of endosomal compartments, while simultaneously blocking autophagic flux by inhibiting fusion of autophagosomes with lysosomes (Mauthe et al., 2018). Monensin treatment induced localization of Gal8, ATG16L1 and LC3 at early endosomes in an ATG13-dependent manner (Fraser et al., 2019), suggesting similarities to lysophagy.

As a morphologically distinctive model of an endosomal compartment, entotic phagosomes or latex bead-containing phagosomes had been analyzed (Florey et al., 2015; Jacquin et al., 2017). Treatment with chloroquine or monensin was found to induce an LAP-like process of direct LC3 recruitment to the intact single phagosomal membrane, without ubiquitination, PI3P production, or WIPI involvement, and independent of ATG13 and of the FIP200-binding domain of ATG16L1.

In the present study, we analyzed the connection between Rabaptin5 and autophagy. We found Rabaptin5 to interact with FIP200 and with the WD domain of ATG16L1 to selectively target autophagy to early endosomes damaged by chloroquine or monensin treatment. In addition, we show that Rabaptin5 initiates autophagy of early *Salmonella*-containing vacuoles and is thus responsible for killing a significant fraction of bacteria early upon infection.

## RESULTS

### FIP200 is a novel interactor of Rabaptin5 on early endosomes

To identify new interaction partners of Rabaptin5, we used human Rabaptin5 as a bait for a yeast two-hybrid screen with a HeLa cell prey library. The screen reproduced the known interactions of Rabaptin5 with itself and with Rabex5, but also revealed new candidates (Suppl. Table S1). Very interestingly, the screen detected FIP200 (Figure 1A and B), a component of the ULK1–ATG13 autophagy initiator complex, as a high confidence interactor of Rabaptin5. The smallest isolated interacting fragment encompassed residues 281–439 of FIP200, a segment outside the coiled-coil regions of the protein (Figure 1C). Importantly, an interaction of FIP200 with Rabaptin5 could be confirmed in HeLa and HEK293A cells by co-immunoprecipitation from cell lysates (Figure 1D).

**Figure 1.**
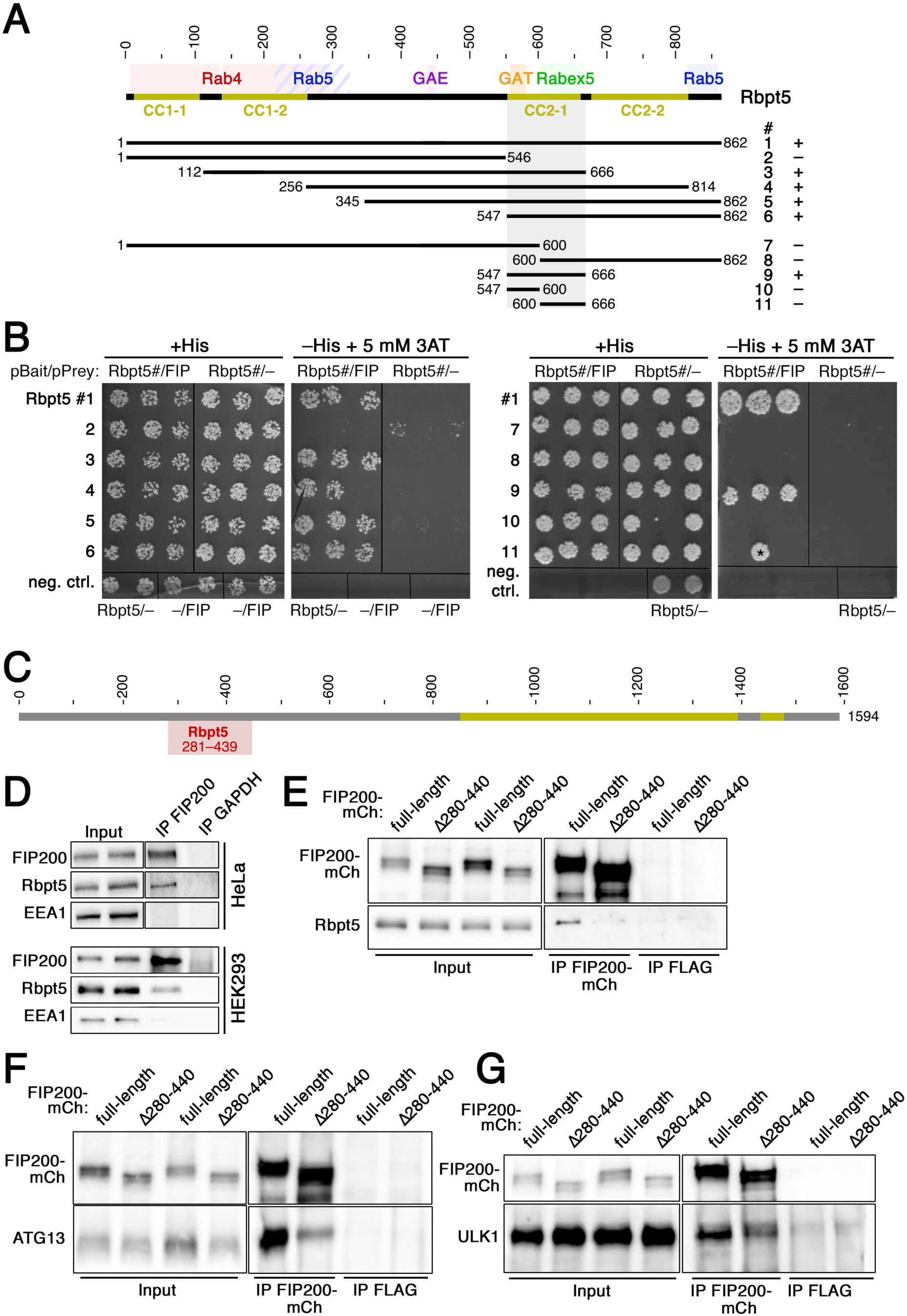
Interaction of Rabaptin5 with FIP200 as identified by two-hybrid analysis in yeast and by co-immunoprecipitation in HeLa and HEK293A cells. **A**: Schematic representation of the sequence of Rabaptin5. Coiled-coil (CC) segments are shown in yellow. Colored backgrounds highlight the segments shown to interact with Rab4, Rab5, Rabex5, and the GAE and GAT domains of GGAs (Golgi-localizing, γ-adaptin ear homology domain, ARF-binding proteins). Below, the segments used to test yeast two-hybrid interaction with residues 257–444 of FIP200 are shown with their number (#) and the observed interaction (+ or –). **B**: Yeast two-hybrid analysis for interaction between the above-shown Rabaptin5 segments (Rbpt5#, fused to LexA on the bait plasmid) and residues 257–444 of FIP200 (FIP, fused to the Gal4 activation domain on the prey plasmid) to drive HIS3 expression. Three different clones each were replica-plated on medium with His or without His, but containing 3-amino-1,2,4-triazole (3AT; an inhibitor of His synthesis to increase stringency) and grown in the absence of Trp and leucine as a control. As negative controls, empty bait or prey plasmids were used. The asterisk indicates a clone invalidated by recombination. **C**: Schematic representation of the sequence of FIP200 with its coiled coil segments in yellow. Residues 281– 439 (gray) indicates the minimal sequence identified to interact with Rabaptin5 in the yeast two-hybrid screen. **D**: FIP200 was immunoprecipitated (IP) from lysates of HeLa or HEK293A cells and probed for FIP200, Rabaptin5 (Rbpt5), and EEA1 (early endosome antigen 1) by immunoblotting. Input lysate (10%) was immunoblotted blotted parallel. As a negative control, the immunoprecipitation was performed using an anti-GAPDH antibody. **E–G**: Lysates of HeLa cells transiently transfected with full-length FIP200-mCherry (FIP200-mCh) or a deletion mutant lacking the segment interacting with Rabaptin5 (Δ280–440) were immuprecipitated with anti-mCherry (IP FIP200-mCh) or, as a control, with anti-FLAG antibodies (IP FLAG). Immunoprecipitates and input lysates (10%) were immunoblotted for mCherry and Rabaptin5 (E), ATG13 (F), or ULK1 (G).

By yeast two-hybrid testing of different protein fragments, the interacting segment in Rabaptin5 was identified to be the coiled-coil domain CC2-1 (Figure 1A and B), where also Rabex5 and the GAT domain of GGA proteins have been shown to bind (Zhang et al., 2014; Zhu et al., 2004). Since this domain and binding of Rabex5 are important for Rabaptin5 recruitment to early endosomes (Kälin et al., 2015), specific disruption of FIP200-binding by its deletion is not possible. Deletion of the segment 280–440 of FIP200 that is sufficient for interaction with Rabaptin5 in the two-hybrid assay was found to be necessary for co-immunoprecipitation of Rabaptin5 (Figure 1E). Unfortunately, deletion of this segment also strongly reduced co-immunoprecipitation of ATG13 and to a lesser extent of ULK1 with FIP200 (Figure 1F and G, resp.), indicating that it did not specifically abrogate binding to Rabaptin5, but also destabilized the incorporation of FIP200 into the ULK1 initiator complex. This is in line with recent structural data indicating that overlapping residues 435–442 are important for protein stability and that residues 443-450 of FIP200, right next to the Rabaptin5-binding region, binds ATG13 (Shi et al., 2020). The FIP200Δ280–440 mutant thus cannot help to define the role of Rabaptin5 in autophagy.

FIP200 transfected alone in HeLa cells localized to small intracellular puncta throughout the cell (Suppl. Figure S1A), corresponding to steady-state autophagosomes (Hara et al., 2008). When expressed together with wild-type Rabaptin5, they colocalized in slightly larger structures (Suppl. Figure S1B). Similarly, FIP200 colocalized with the co-transfected early endosomal markers Rab4 and Rab5, but not with the late endosomal marker Rab7 (Suppl. Figure S1C–E). This observation is consistent with an interaction of FIP200 with early Rabaptin5-positive endosomes.

### Chloroquine treatment induces autophagy of Rabaptin5-positive early endosomes

To assess a potential involvement of Rabaptin5 in autophagy at early endosomes, we employed chloroquine treatment as in previous studies (e.g. Mauthe et al., 2018). To be able to easily detect Rabaptin5-positive early endosomes by immunofluorescence without transient transfection, we generated a stable HEK293A cell line moderately overexpressing Rabaptin5 approximately 5 times (HEK^+Rbpt5^; Figure 2A and A’). Incubation of these cells with 60 µM chloroquine for 30 min indeed produced swollen structures appearing as small rings positive for Rabaptin5 and transferrin receptor and thus identifiable as early endosomes (Figure 2B). In transfected cells expressing moderate levels of mCherry-galectin3, this lectin was found to be recruited to the Rabaptin5-positive endosomes (Figure 2C), indicating membrane rupture. They also stained positive for ubiquitin (Figure 2D), consistent with exposure of lumenal protein domains to the cytosol. These rings further stained positive for early autophagy components. Transfected mCherry-FIP200 strongly colocalized with Rabaptin5-positive structures (Figure 2E), which appeared enlarged most likely due to the overexpression of FIP200. WIPI2 and ATG16L1 (Figure 2F and G) also localized to these early endosomes. At this early time point of 30 min, the late component LC3B had not accumulated yet (Figure 2H).

**Figure 2.**
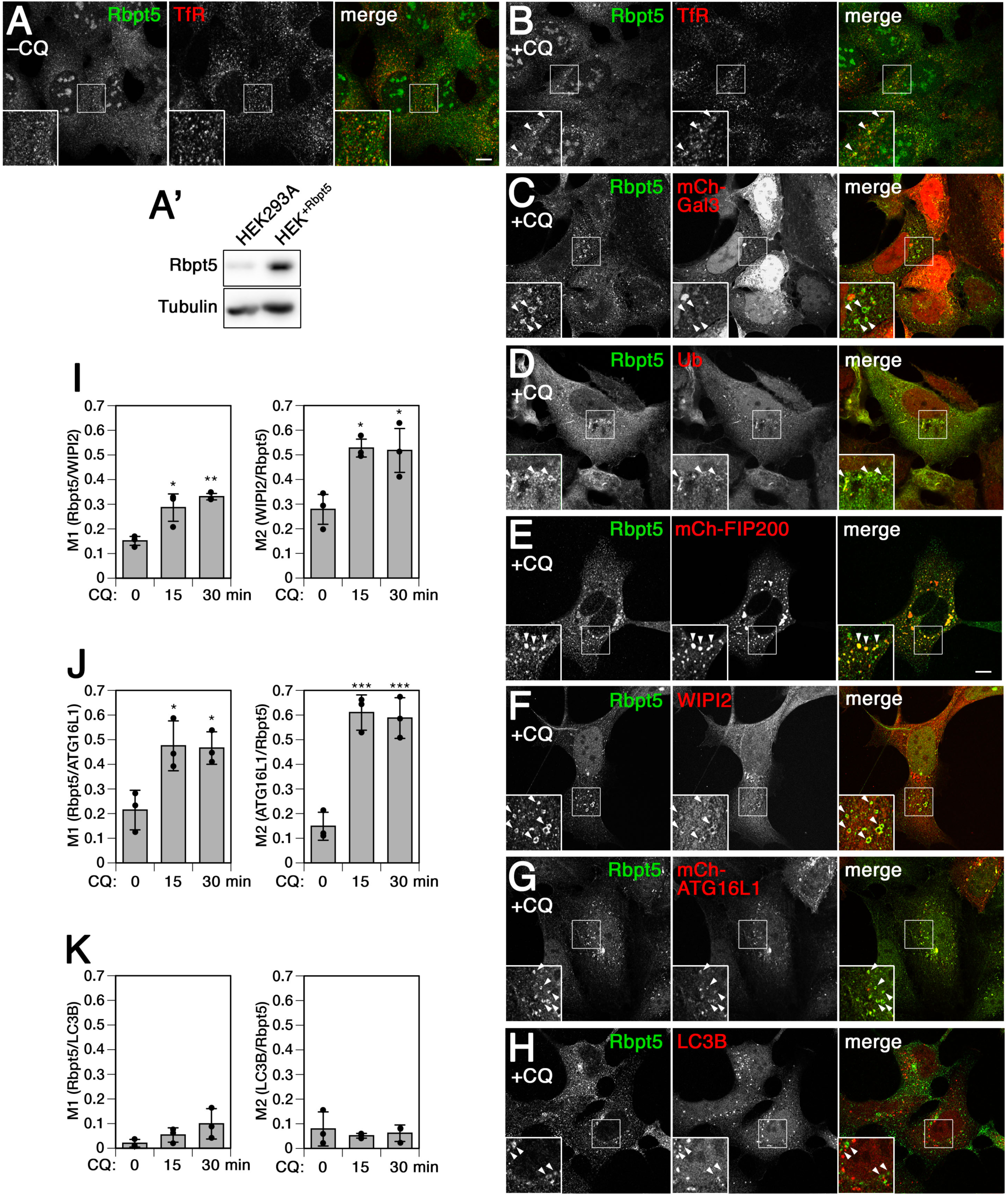
Autophagy proteins are targeted to Rabaptin5-positive endosomes upon chloroquine treatment. **A–H**: To more easily visualize Rabaptin5, a stable HEK293A cell line overexpressing Rabaptin5 (HEK^+Rbpt5^) was generated. Rabaptin5 levels were anayzed by immunoblotting in comparison to parental HEK293A cells (panel **A’**). HEK^+Rbpt5^ cells, untransfected or 24 h after transfection with mCherry-Galectin3, mCherry-FIP200, or mCherry-ATG16L1, without (–CQ) or with chloroquine treatment (60 µM) for 30 min (+CQ), were analyzed by immunofluorescence microscopy for Rabaptin5 and either transferrin receptor (TfR) (**A** and **B**), mCherry-Galectin3 (**C**), ubiquitin (**D**), mCherry-FIP200 (**E**), WIPI2 (**E**), mCherry-ATG16L1 (**G**), or LC3B (**H**). Bar, 10 µm. In the enlarged insets, arrowheads point out chloroquine-induced enlarged (circular) early endosomes. **I–K**: HEK^+Rbpt5^ cells, untransfected or 24 h after transfection with mCherry-ATG16L1, were treated with 60 µM chloroquine for 0, 15 and 30 min and stained for Rabaptin5 and either WIPI2, mCherry-ATG16L1, or LC3B. Manders’ colocalization coefficients were determined, M1 showing the fraction of Rabaptin5-positive structures also positive for WIPI2 (**I**), mCherry-ATG16L1 (**J**), or LC3B (**H**), and M2 showing the respective inverse (mean and standard deviation of three independent experiments; ANOVA: *p < 0.05, **p < 0.01, ***p<0.001).

Colocalization of WIPI2 and ATG16L1 was not only observed qualitatively on large Rabaptin5-positive rings, but also globally as quantified using Manders’ colocalization coefficients (Figure 2I–K): the fraction of Rabaptin5 on WIPI2- or ATG16L1-positive structures (M1) and the fractions of these two proteins on Rabaptin5-positive structures (M2) were significantly increased already after 15 min chloroquine treatment and similarly after 30 min, while no significant colocalization was observed with LC3B.

A rapid effect of endomembrane damage has been reported to be membrane repair by an ESCRT-mediated mechanism involving ALIX and the ESCRT-III effector CHMP4B on damaged lysosomes (Jia et al., 2020; Radulovic et al., 2018; Skowyra et al., 2018). Lysosome disruption and subsequent lysophagy can be specifically induced by L-leucyl-L-leucine methyl ester (LLOMe) that is condensed into a membranolytic polymer by cathepsin C in lysosomes (Chauhan et al., 2016; Maejima et al., 2013). While 30-min incubation with LLOMe strongly recruited the CHMP4B and ALIX to damaged lysosomes, chloroquine treatment caused hardly any activation of CHMP4B or ALIX (Suppl. Figure S2). No recruitment of either ESCRT protein was observed to enlarged Rabaptin5-positive endosomes.

### Chloroquine-induced autophagy, but not Torin1-induced autophagy or lysophagy, depends on Rabaptin5

Upon longer exposure of 150 min to chloroquine or to Torin1, an inhibitor of mTOR to induce autophagy mimicking starvation, control cells showed a strong increase in the numbers of both early WIPI2-positive and late LC3B-positive autophagosomal puncta (Figure 3A–C). Silencing of FIP200 expression by RNA interference blocked this increase of WIPI2 autophagosomes completely and of LC3B autophagosomes to a large extent for both treatments. Silencing of Rabaptin5 similarly inhibited specifically the formation of chloroquine-induced WIPI2 and LC3B structures, but did not affect autophagy induction by Torin1. This result indicates that Rabaptin5 is specifically involved in autophagy triggered by chloroquine, but not in starvation-induced autophagy regulated by mTOR, while FIP200 is required for both. The observation that knockdown of Rabaptin5 as well as of FIP200 appears less effective in reducing LC3B structures with chloroquine than WIPI2 puncta is likely due to stabilization of late autophagosomes by inhibition of autophagic flux (e.g. Mauthe et al., 2018).

**Figure 3.**
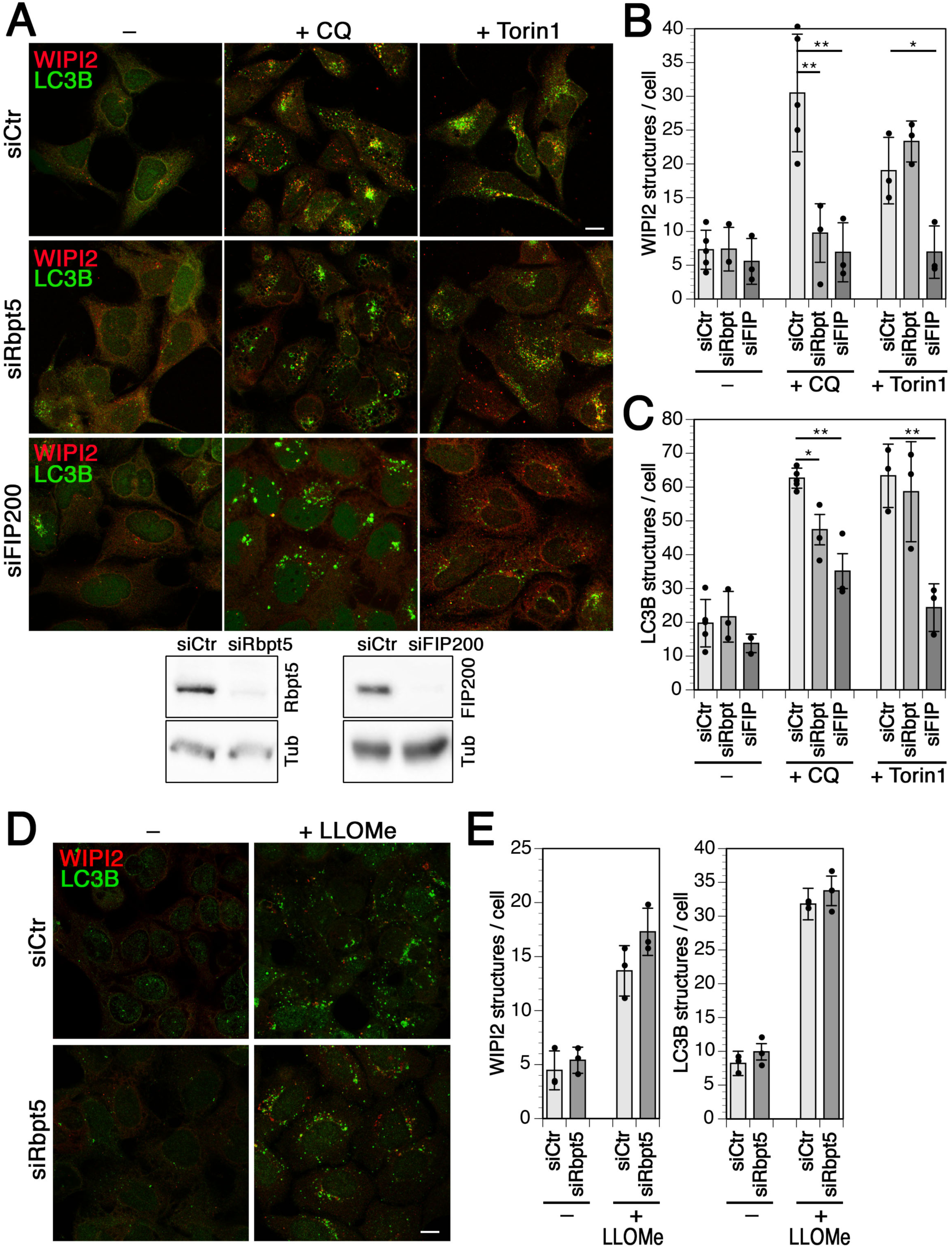
Chloroquine-, but not Torin1- or LLOMe-induced autophagy depends on Rabaptin5. **A**: HEK293A cells were transfected with nontargeting siRNA (siCtr) or siRNAs silencing Rabaptin5 (siRbpt5) or FIP200 (siFIP200) for 72 h and treated without (–) or with 60 µM chloroquine (+CQ) or 250 nM Torin1 for 150 min. Below, the efficiency of Rabaptin5 and FIP200 knockdown was assayed by immunoblotting using tubulin (Tub) as a loading control. **B and C**: WIPI2 (B) or LC3B puncta per cell (C) were quantified for each condition (mean and standard deviation of three independent experiments; ANOVA: *p < 0.05, **p < 0.01). **D**: HEK293A cells were transfected with siCtr or siRbpt5 as in A and incubated without or with 280 µM LLOMe for 150 min to induce lysophagy. Cells were fixed and immunostained for endogenous WIPI2 and LC3B. Bar, 10 µm. **E**: WIPI2 or LC3B puncta per cell were quantified (mean and standard deviation of three independent experiments).

Since chloroquine more generally affects acidified organelles, we attempted to differentiate between autophagy of lysosomes (lysophagy) and of endosomes. Incubation with LLOMe for 150 min induced the formation of WIPI2 and LC3B puncta that was not affected by silencing of Rabaptin5 (Figure 3D and E), indicating a specific role of Rabaptin5 in autopagy of damaged endosomes.

### Chloroquine-induced autophagy requires the FIP200 complex and ATG16L1

We further tested for an involvement of ULK1 and ATG13, the two main partners of FIP200 in the classical autophagy initiation complex, in chloroquine-induced autophagy (Figure 4A and B). Silencing of ATG13 essentially blocked an increase of WIPI2 puncta and strongly reduced LC3B structures, just like silencing of FIP200 or Rabaptin5 did. In contrast, silencing of ULK1, which inhibited starvation- and Torin1-induced autophagy (Suppl. Figure S3), did not significantly reduce chloroquine induction of WIPI2- or LC3B-positive autophagosomes (Figure 4A and B). To test whether ULK2 might replace its homolog ULK1 in this process, we used the general ULK inhibitor MRT68921 (Chaikuad et al., 2019; Petherick et al., 2015). Incubation of cells with MRT68921 entirely blocked induction of WIPI2 or LC3B autophagosomes by chloroquine (Figure 4C and D), indicating that ULK2 or either one of the two ULK proteins is required for this function. FIP200 thus acts in chloroquine-induction of autophagy together with its complex partners.

**Figure 4.**
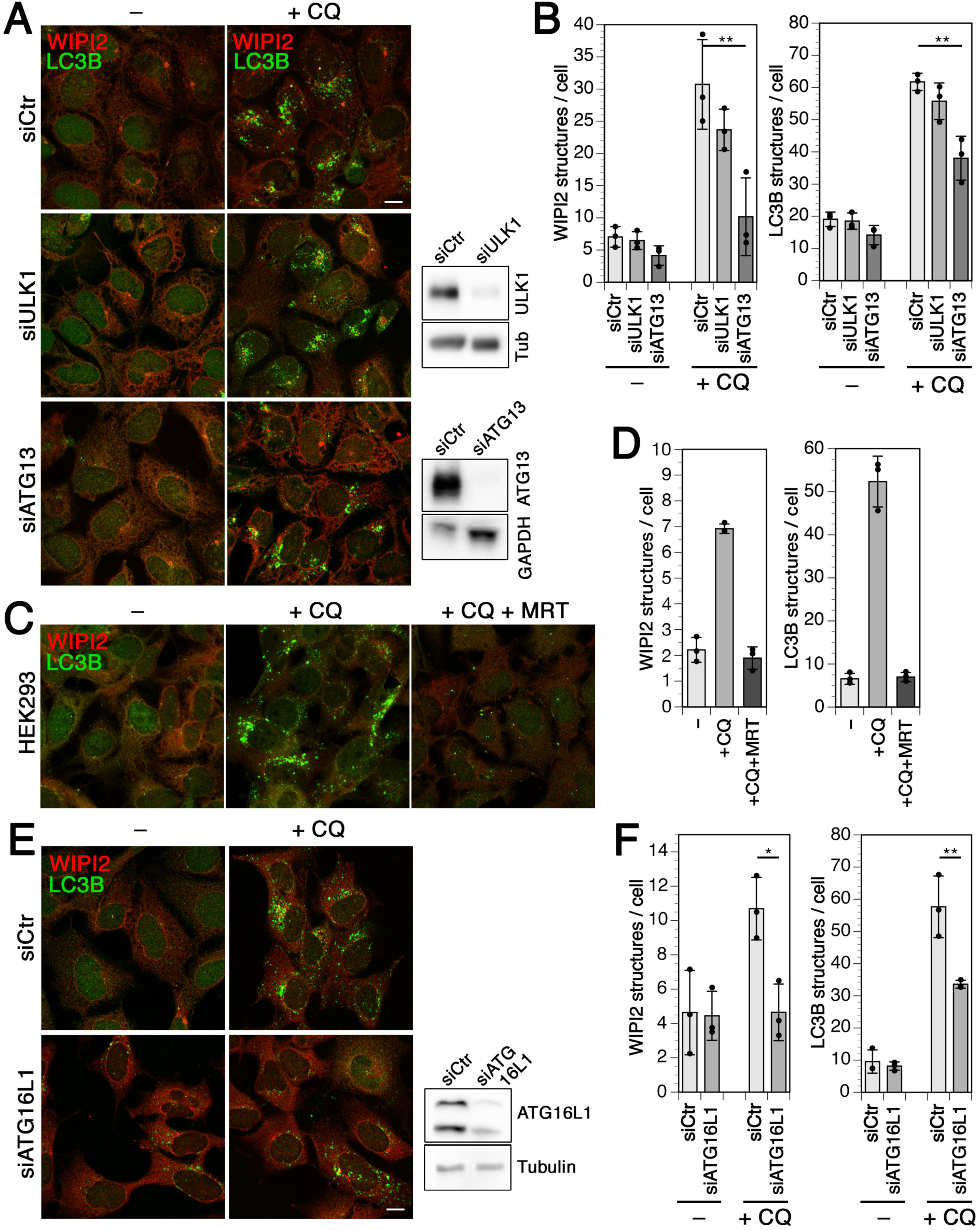
Chloroquine-induced autophagy requires the FIP200–ULK1/2–ATG13 complex. **A**: HEK293A cells were transfected with nontargeting siRNA (siCtr) or siRNAs silencing ULK1 (siULK1) or ATG13 (siATG13) for 72 h, treated without (–) or with 60 µM chloroquine (+CQ) for 150 minutes, and fixed and immunostained for endogenous WIPI2 and LC3B. Bar, 10 µm. Efficiency of ULK1 and ATG13 knockdown was assayed by immunoblotting on the right using tubulin (Tub) or GAPDH as loading controls. **B**: WIPI2 of LC3B puncta per cell were quantified for each condition (mean and standard deviation of three independent experiments; ANOVA: *p < 0.05, **p < 0.01). **C**: HEK293A cells were incubated without (–), with 60 µM chloroquine (+CQ), or with chloroquine and 5 µM MRT68921 (+CQ +MRT) for 150 min and fixed and stained for WIPI2 and LC3B. Bar, 10 µm. **D**: WIPI2 of LC3B puncta per cell were quantified for each condition as in panel B. **E**: Cells were transfected with control siRNA or siRNA silencing ATG16L1, treated for 150 min with chloroquine as in (A), and fixed and stained for WIPI2 and LC3B. Bar, 10 µm. Efficiency of ATG16L1 knockdown was assayed by immunoblotting on the right. **F**: WIPI2 of LC3B puncta per cell were quantified for each condition as in panel B.

Silencing of ATG16L1, which recruits the ATG5/12 E3-like enzyme complex for LC3 lipidation, not only reduced LC3B autophagosomes induced by chloroquine, but also blocked formation of WIPI2 puncta (Figure 4C and D). ATG16L1 thus appears to be required to recruit or stabilize WIPI2. It has already been reported that WIPI2 puncta were reduced in cells lacking ATG16L1 or expressing ATG16L1 mutated in its PI3P-binding motif (Dudley et al., 2019). But as expected from previous studies (Bakula et al., 2017; Dooley et al., 2014; Dudley et al., 2019), the opposite was also the case: recruitment of mCherry-ATG16L1 to chloroquine-induced enlarged Rabaptin5-positive endosomes as well as colocalization in general was inhibited upon siRNA silencing of WIPI2 (Suppl. Figure S4A). This indicates mutual stabilization of WIPI2 and ATG16L1 on damaged early endosomes.

### Rabaptin5 binds to the WD domain of ATG16L1 via a conserved interaction motif

To test whether Rabaptin5 interacts with ATG16L1 on early endosomes, similar to TRIM16 or TMEM59 in autophagy of lysosomes or bacteria-containing phagosomes (Boada-Romero et al., 2013; Chauhan et al., 2016; Jia et al., 2020), we performed co-immunoprecipitation experiments. Indeed, ATG16L1 was co-immunoprecipitated with Rabaptin5 at low levels already from untreated HeLa cells and more extensively after 30 min of chloroquine treatment (Figure 5A). The interaction decreased after 2 h with chloroquine. Co-immunoprecipitation was also observed in HEK293A cells after a 30-min incubation with chloroquine (Figure 5B). However, in a cell line lacking FIP200 generated by CRISPR/Cas9 gene inactivation, no specific co-isolation with immunoprecipitated Rabaptin5 was detected (Figure 5B). In FIP200 knockout cells, mCherry-ATG16L1 recruitment to chloroquine-induced enlarged Rabaptin5-positive endosomes and colocalization between Rabaptin5 and mCherry-ATG16L1 were inhibited (Suppl. Figure S4B). Endosomal autophagy and interaction of Rabaptin5 with ATG16L1 thus depend on FIP200.

**Figure 5.**
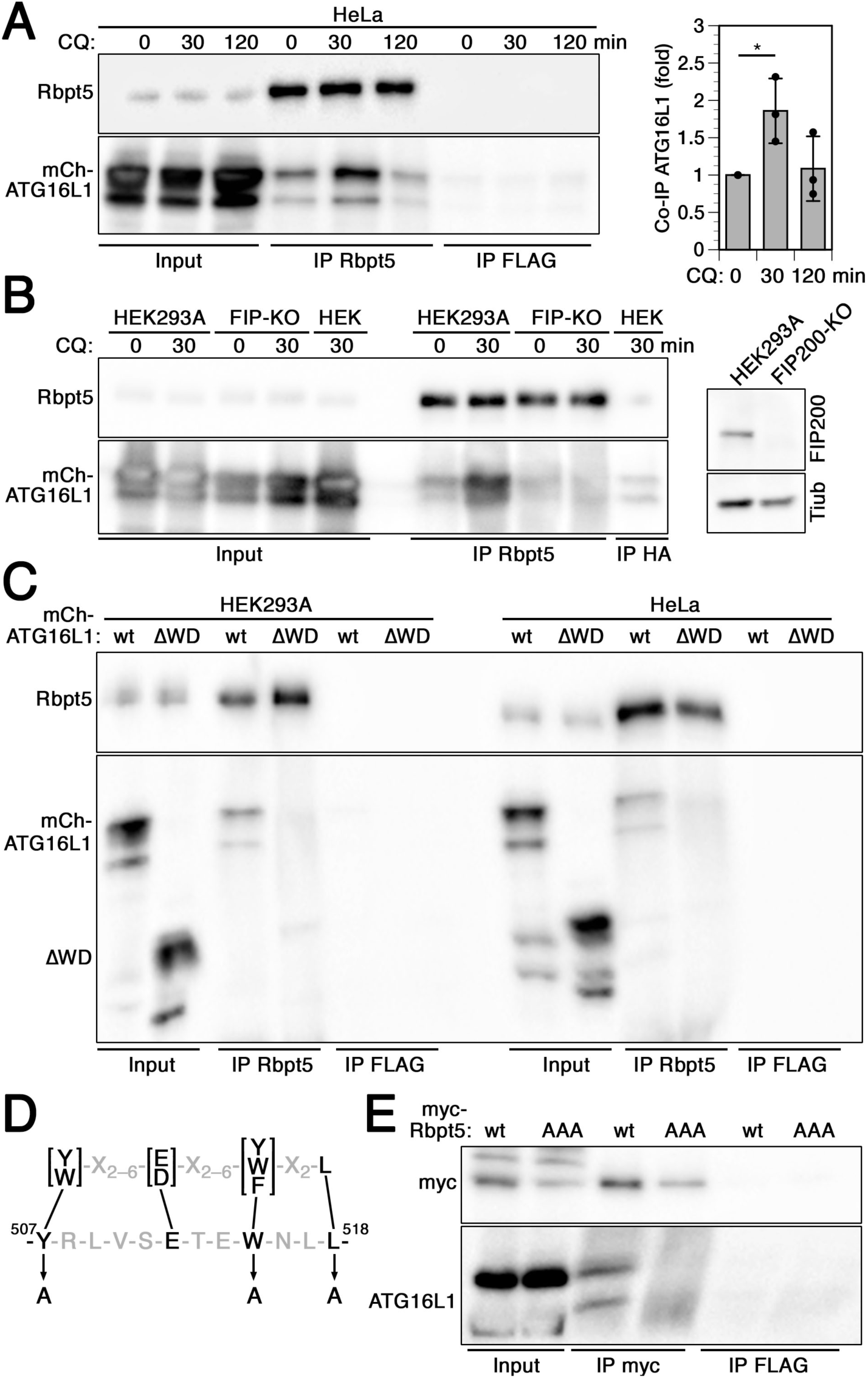
Rabaptin5 binds to the WD domain of ATG16L1 via a conserved interaction motif. **A**: HeLa cells transiently transfected with full-length mCherry-ATG16L1 were treated with 60 µM chloroquine for 0, 30, or 120 min, lysed, and immunoprecipitated with anti-Rabaptin5 (IP: Rbpt5) or, as a control, with anti-FLAG antibodies (IP FLAG). Immunoprecipitates and input lysates (10%) were immunoblotted for Rabaptin5 and ATG16L1. Signals were quantified and the ratios of mCherry-ATG16L1/Rabaptin5 normalized to that without (0 min) chloroquine treatment (mean and standard deviation of three independent experiments; ANOVA: *p < 0.05). **B**: Co-immunoprecipitation was performed as in panel A using parental HEK293A cells and CRISPR/Cas9 knockout cells lacking FIP200 (FIP-KO). Anti-HA antibodies were used as a control (IP HA). On the right, HEK293A and FIP200 knockout cells were immunoblotted for FIP200 and as a loading control of tubulin (Tub). **C**: Lysates of HEK293A or HeLa cells transiently transfected with full-length mCherry-ATG16L1 (wt) or a mutant lacking the WD domain (ΔWD) were immunoprecipitated with anti-Rabaptin5 or anti-FLAG antibodies, and immunoblotted for Rabaptin5 and ATG16L1. **D**: The consensus sequence of the ATG16L1 interaction motifs of TMEM59, NOD2, and TLR2 (above; Boada-Romero et al. 2013) is shown together with the matching sequence in Rabaptin5 (below). The three point mutations to alanine to produce the AAA mutant of Rabaptin5 are indicated. **E**: Lysates of HeLa cells transiently transfected with myc-tagged wild-type Rabaptin5 (wt) or triple-alanine mutant (AAA) were immunoprecipitated with anti-myc (IP myc) or anti-FLAG antibodies (IP FLAG), and immunoblotted for myc and ATG16L1.

Co-immunoprecipitation of ATG16L1 with Rabaptin5 was lost upon deletion of the WD-repeat domain in ATG16L1 (Figure 5C). Rabaptin5 thus interacts differently with ATG16L1 than TRIM16, which was shown to bind to the coiled-coil domain (Chauhan et al., 2016). In contrast, the membrane protein TMEM59, a promoter of selective autophagy of *Staphylococcus aureus*-containing phagosomes, had previously been shown to bind to the WD domain (Boada-Romero et al., 2013). A 19-amino acid peptide motif containing four essential residues had been identified to mediate this interaction. Similar largely conserved motifs were also found in NOD2, a protein known to recruit ATG16L1 at bacterial entry sites, and in TLR2 that promotes LC3 lipidation at phagosomes. Rabaptin5 also contains a sequence (residues 507–518) that complies with this consensus motif (Figure 5D). Mutation of the three most important residues of this motif to alanines (Y507A/W515A/L518A in Rabaptin5AAA), which in TMEM59 completely inactivated ATG16L1 binding, also eliminated co-immunoprecipitation of ATG16L1 with mutant Rabaptin5 (Figure 5E).

### Binding of Rabaptin5 to ATG16L1 is essential for endosomal autophagy

To assess the importance of the interaction between Rabaptin5 and ATG16L1, we produced a Rabaptin5 knockout HEK293A cell line (Rbpt5-KO) by CRISPR/Cas9 gene inactivation. By transfecting wild-type Rabaptin5 or the triple mutant Rabaptin5-AAA, we produced the stable cell lines Rbpt5-KO+wt and Rbpt5-KO+AAA that produced similar Rabaptin5 levels as the original HEK293A cell line (Figure 6A). Again, we analyzed WIPI2- and LC3-positive autophagosomes before and after chloroquine treatment for 150 min (Figure 6B and C). While overexpression of Rabaptin5 resulted in a slight increase of steady-state and chloroquine-induced WIPI2 puncta, Rabaptin5 knockout left the steady-state levels unchanged and eliminated chloroquine-induced formation of WIPI2 autophagosomes (Figure 6C, left). Re-expression of wild-type Rabaptin5 rescued WIPI2 autophagosome induction by chloroquine, while Rabaptin5-AAA did not. This result suggests that the interaction of Rabaptin5 with ATG16L1 is essential for early endosomal autophagy upon chloroquine damage.

**Figure 6.**
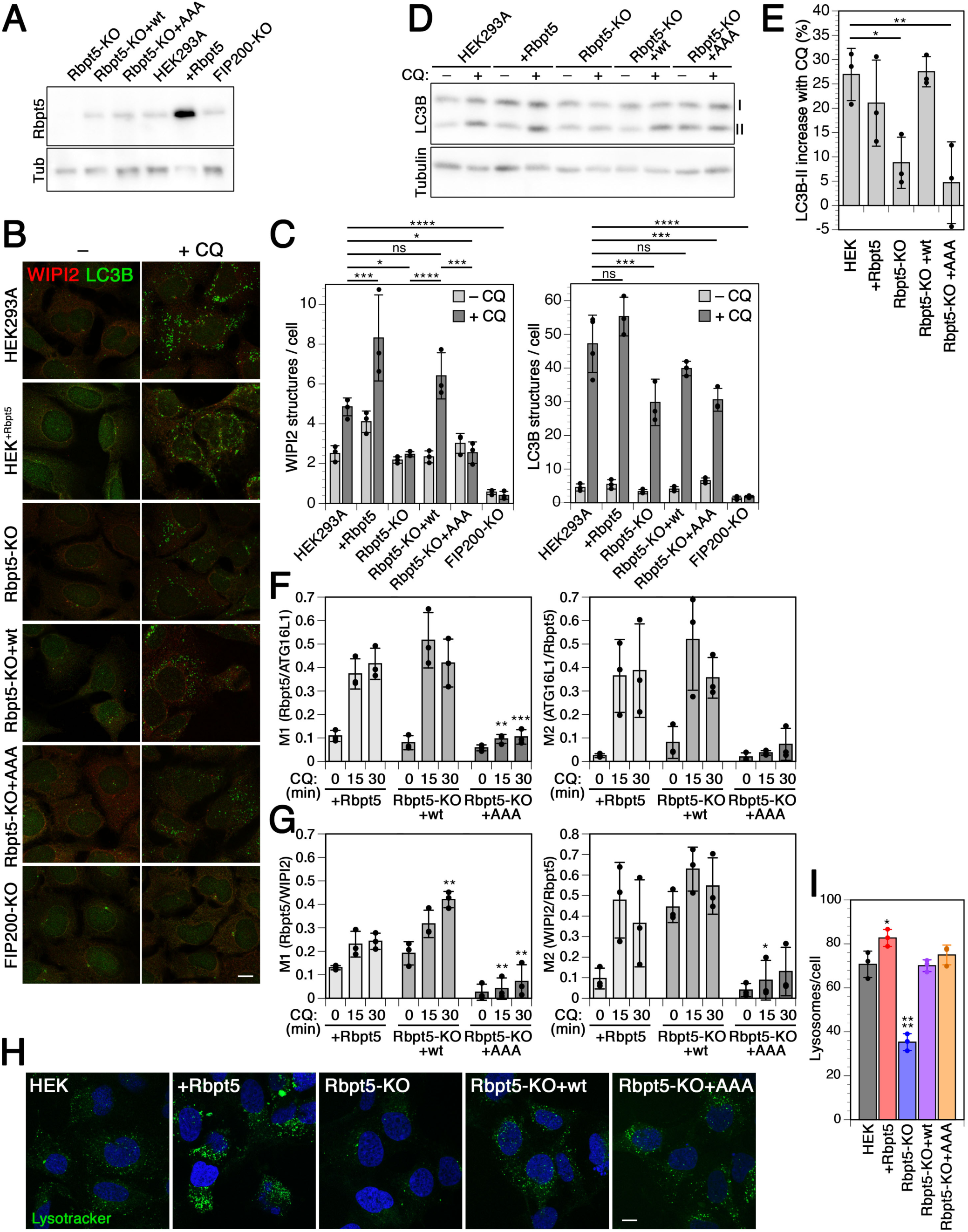
Chloroquine-induced endosomal autophagy depends on Rabaptin5 and its ATG16L1 binding motif. **A**: By immunoblot analysis, the levels of Rabaptin5 and as a loading control of tubulin (Tub) were assessed in wild-type HEK293A cells, HEK^+Rbpt5^ cells stably overexpressing Rabaptin5, and Rabaptin5-knockout cells without (Rbpt5-KO) or with stable re-expression of wild-type (Rbpt5-KO+wt) or AAA-mutant Rabaptin5 (Rbpt5-KO+AAA). **B**: The same stable HEK293A-derived cell lines were treated without (–CQ) or with 60 µM chloroquine for 150 min (+CQ) and analyzed by immunofluorescence microscopy for WIPI2 or LC3B. **C**: WIPI2 and LC3B puncta per cell were quantified for each condition (mean and standard deviation of three independent experiments; ANOVA: *p < 0.05, **p < 0.01, ***p < 0.001, ****p < 0.0001). **D and E:** The HEK293A-derived cell lines were treated with 60 µM chloroquine for 30min and non-lipidated and lipidated LC3B (I and II, resp.) were assayed by immunoblot analysis (D). The increase of the LC3B-II fraction of total LC3B upon chloroquine treatment was quantified (mean and standard deviation of three independent experiments; ANOVA: *p < 0.05, **p < 0.01). **F**: HEK^+Rbpt5^ cells and Rabaptin5-knockout cells stably re-expressing wild-type (Rbpt5-KO+wt) or AAA-mutant Rabaptin5 (Rbpt5-KO+AAA) were transfected with mCherry-ATG16L1, treated with 60 µM chloroquine (CQ) for 0, 15 and 30 min, and analyzed by immunofluorescence microscopy for Rabaptin5 and mCherry-ATG16L1. Manders’ colocalization coefficients were determined, M1 showing the fraction of Rabaptin5-positive structures also positive for mCherry-ATG16L1 and M2 showing the inverse (mean and standard deviation of three independent experiments; ANOVA for Rbpt5-KO+wt or +AAA vs. HEK^+Rbpt5^ cells: *p < 0.05, **p < 0.01, ***p < 0.001). **G**: The same three cell lines as in panel F, but untransfected, were analyzed in the same way for Rabaptin5 and WIPI2. **H**: Wild-type HEK293A cells, HEK^+Rbpt5^ cells, Rabaptin5-knockout cells without (Rbpt5-KO) or with stable re-expression of wild-type (Rbpt5-KO+wt) or AAA-mutant Rabaptin5 (Rbpt5-KO+AAA) were stained with lysotracker and with DAPI for nuclei. Bar, 10 µm. **I**: The number of lysosomes (lysotracker-positive structures) per cell was quantified from cells as in panel C (mean and standard deviation of three independent experiments). Images for at least 15 cells per sample were quantified. The value for HEK^+Rbpt5^ cells is an underestimate, because the density of puncta makes them difficult to distinguish as separate structures. ANOVA: *p < 0.05, ****p < 0.0001.

Rabaptin5 knockout also reduced the chloroquine-dependent increase of LC3B-positive puncta (Figure 6C, right), consistent with inhibition of endosomal autophagy, but did not block it. Again, re-expression of wild-type Rabaptin5, but not Rabaptin5-AAA, rescued formation of LC3B-positive structures. FIP200 knockout strongly reduced both steady-state and chloroquine-induced autophagosomes positive for either WIPI2 or LC3B, consistent with an essential role for both steady-state and endosomal autophagy.

These results with stable cell lines confirm those with siRNA knockdown of Rabaptin5 or FIP200 of Figure 3. It should be noted that the numbers of WIPI2 and LC3B puncta were consistently higher under knockdown conditions even at steady-state. A direct comparison of untreated and control siRNA transfected cells showed this to be nonspecifically caused by siRNA transfection, reflected in higher numbers of autophagosomes and an increased ratio of lipidated / unlipidated LC3B (Suppl. Figure S5).

Immunoblot analysis of LC3B lipidation upon chloroquine treatment (Figure 6D and E) showed the expected increase of LC3B-II in wild-type and Rabaptin5-overexpressing cells already after 30 min, which was reduced in Rabaptin5 knockout cells and rescued by re-expression of wild-type Rabaptin5, but not of the AAA mutant.

We furthermore quantified colocalization of Rabaptin5 and ATG16L1 or WIPI2 upon chloroquine treatment for 0, 15, or 30 min in Rabaptin5-overexpressing cells and in knockout cells re-expressing wild-type or AAA mutant Rabaptin5 (Figure 6F and G). Colocalization with either autophagy protein was lower and hardly induced in knockout cells expressing the Rabaptin5-AAA mutant unable to bind ATG16L1, demonstrating that this interaction is required to induce endosomal autophagy.

There are a number of interconnections between autophagy and endosomes (Birgisdottir and Johansen, 2020). To address the possibility that absence or mutation of Rabaptin5 might alter endosome properties in a way that indirectly affects chloroquine-induced autophagy, we confirmed that chloroquine induces early endosome swelling also in Rabaptin5-KO cells by immunofluorescence microscopy of EEA1 (Suppl. Figure 6A). Rabaptin5 deletion did not affect autophagic flux, since expression of tandem fluorescent tagged RFP-GFP-LC3 containing an acid-sensitive GFP and a pH-insensitive RFP (Kimura et al., 2007) showed an indistinguishable fraction of autophagosomes (red/green double-positive LC3 puncta) and autolysosomes (red-only LC3 puncta) in HEK293A and Rabaptin5-KO cells, without treatment or upon induction of autophagy by starvation or Torin1 treatment (Suppl. Figure 6B and C).

To further exclude the possibility that the triple AAA mutation inactivated the function of Rabaptin5 in endosomal maturation and thus perhaps affected endosomal autophagy indirectly, we analyzed the number of lysosomal structures in the different HEK cell lines by lysotracker staining (Figure 6H). The marked reduction of lysosomes upon Rabaptin5 knockout was completely rescued by re-expression of wild-type Rabaptin5 and also by Rabaptin5-AAA, demonstrating that the endosomal function of the mutant protein was intact (Figure 6I).

### Monensin-induced endosomal autophagy also depends on Rabaptin5 and its interaction with ATG16L1

Chloroquine is protonated and trapped within acidified endosomes and lysosomes, osmotically promoting compartment swelling by water influx. Monensin promotes the exchange of protons for osmotically active monovalent cations like Na^+^ and thus results in osmotic swelling of acidified compartments in a different way. Monensin treatment of HEK^+Rbpt5^ cells rapidly produced swollen Rabaptin5-positive early endosomes that stained positive for ATG16L1 and WIPI2 (Suppl. Figure S7A), just like chloroquine treatment (Figure 2F and G). Similarly, overall colocalization of ATG16L1 and WIPI2 with Rabaptin5-positive structures increased in Rabaptin5 overexpressing HEK^+Rbpt5^ cells and in Rabaptin5-knockout cells re-expressing wild-type Rabaptin5, but not in knockout cells expressing the Rabaptin5-AAA mutant that is unable to bind to ATG16L1 (Suppl. Figure S7B and C). The increase in total WIPI2-positive structures upon longer monensin treatment of 150 min was also dependent on Rabaptin5, since it was abolished upon knockout of Rabaptin5 and recovered upon re-expression of wild-type Rabaptin5, but not by the AAA mutant (Suppl. Figure S7D). The effect on the induction of LC3B-positive structures was less clearly dependent on Rabaptin5 (Suppl. Figure S7E).

### Rabaptin5 targets *Salmonella*-containing vacuoles to autophagy

As *Salmonella enterica* enters host cells like fibroblasts or epithelial cells, its surrounding phagosomal membrane quickly acquires the characteristics of early endosomes through fusion with Rab5-positive endosomes (LaRock et al., 2015; Levin et al., 2016; Steele-Mortimer et al., 1999). The *Salmonella*-containing vacuole (SCV) is thus positive for Rab5, Rabaptin5/Rabex5, EEA1, transferrin receptor (TfR), and the PI-3 kinase VPS34. Perforation of the early SCV membrane by the Type III secretion system 1 (T3SS-1) results in binding of cytosolic galectins and ubiquitination to activate antibacterial autophagy to the SCV membrane (LaRock et al., 2015). Because of this similarity of the early SCV with damaged early endosomes, we tested the role of Rabaptin5 in autophagy of *Salmonella* early after infection.

We first infected HeLa cells with *Salmonella enterica* serovar Typhimurium with or without siRNA-mediated silencing of Rabaptin5 or FIP200 and determined the number of phagocytosed live bacteria after 0, 1, 3 and 6 h (Figure 7A). For this, bacteria were gently centrifuged onto the cell layer for 5 min at 37°C and further incubated for 10 min. Bacteria that had not entered the cells were thoroughly washed away. For further incubation, the medium was supplemented with gentamycin to eliminate any remaining extracellular bacteria. Yet, even immediately after infection, confocal immunofluorescence microscopy confirmed that essentially all cell-associated bacteria were internalized. The infection efficiency was not affected by silencing Rabaptin5 or FIP200 (∼14% of cells infected; Suppl. Figure S8). At different time points, cells were lysed and plated out to determine the number of live bacteria.

**Figure 7.**
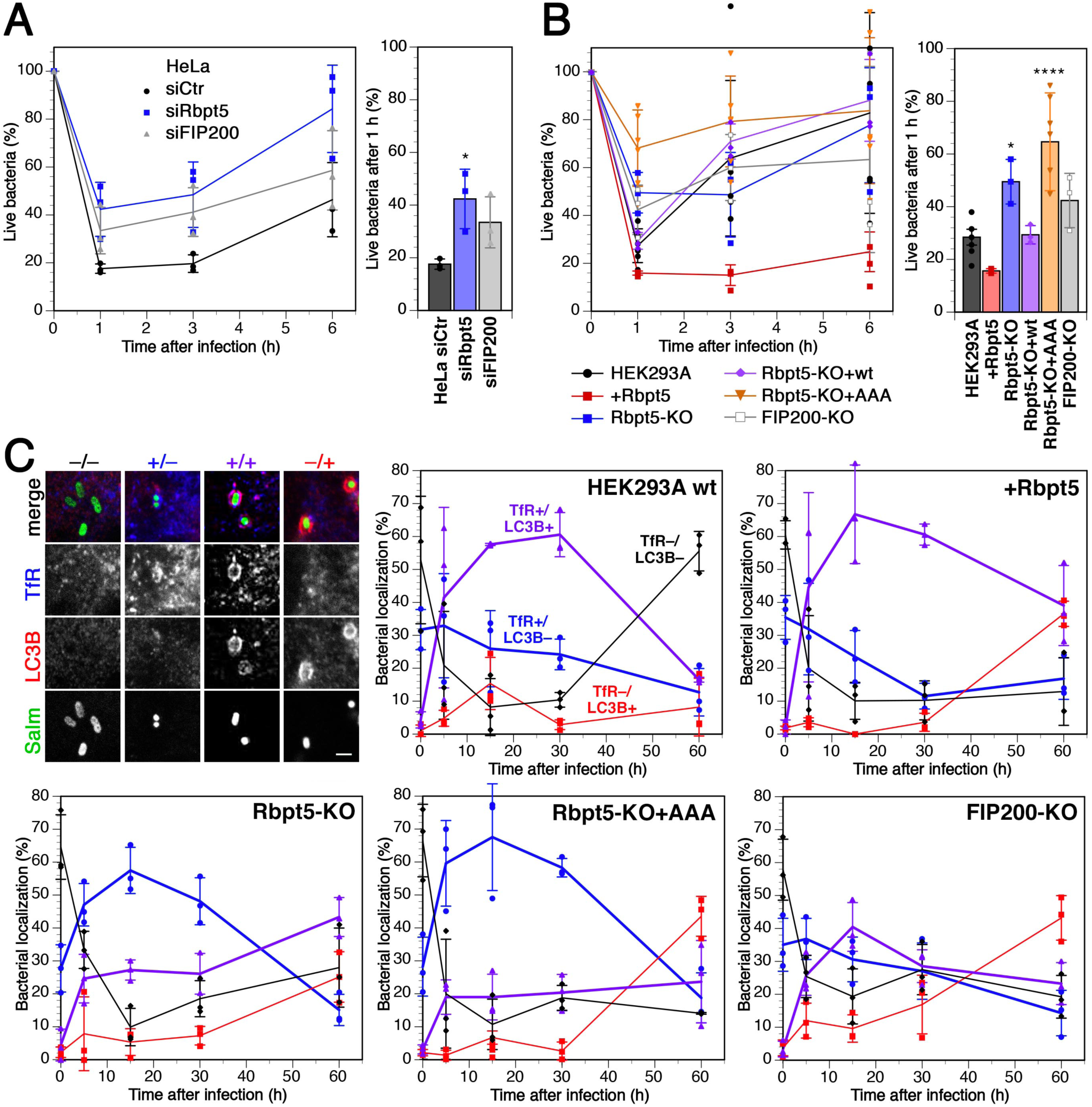
Rabaptin5-mediated autophagy contributes to early killing of *Salmonella*. **A**: HeLa cells were transfected with nontargeting siRNA (siCtr) or siRNAs silencing Rabaptin5 (siRbpt5) or FIP200 (siFIP200) for 72 h. The cells were infected with *Salmonella* by centrifugation at 500×g for 5min at 37°C and incubation for 10 min at 37°C, washed three times, and incubated in fresh culture medium containing gentamicin to prevent growth of extracellular bacteria for 0, 1, 3, or 6 h before lysis of the host cells and plating of the bacteria on LB agar plates at various dilutions to determine the number of live bacteria at the different time points, shown as a percentage of internalized cells after infection (mean and standard deviation of three independent experiments). On the right, the fractions of internalized bacteria alive 1 h after infection are shown separately (mean and standard deviation of three independent experiments; ANOVA: *p < 0.05). **B**: Wild-type HEK293A, HEK^+Rbpt5^, Rbpt5-KO, Rbpt5-KO+wt, Rbpt5-KO+AAA, and FIP200-KO cells were infected with *Salmonella* and treated and analyzed as in panels A (mean and standard deviation of three independent experiments; ANOVA: *p < 0.05, ****p < 0.0001). **C**: Wild-type HEK293A, HEK^+Rbpt5^, Rbpt5-KO, Rbpt5-KO+AAA, and FIP200-KO cells were infected with *Salmonella* expressing GFP as in panel B, incubated in fresh culture medium containing gentamicin for 0, 5, 15, 30, and 60 min, fixed with methanol and immunostained for transferrin receptor (TfR) as a marker of early endosomes and for LC3B as a marker of autophagy. *Salmonella* were classified according to their association with a TfR- and/or LC3B-positive compartment – as illustrated on the top left (bar, 2 µm) – during the first hour after infection. In the absence of Rabaptin5, LC3-positive SCVs with early endosomal characteristics (containing TfR) were strongly reduced. (Means and standard deviations of three independent experiments, analyzing >50 bacteria for each time point.)

Within the first hour of infection, ∼80% of initially internalized bacteria were killed, after which live cell numbers increased again. Part of the observed initial killing is due to autophagy, since knockdown of FIP200 almost doubled the number of live cells 1 h after infection (Figure 7A, right). Silencing of Rabaptin5 expression was at least as effective, indicating a contribution of Rabaptin5 to early bacterial killing. These findings were confirmed by experiments using our stable HEK293A cell lines (Figure 7B). Only 25% of internalized *Salmonella* survived the first hour in parental HEK293A cells, whereas overexpression of Rabaptin5 in HEK^+Rbpt5^ cells reduced this fraction by half and knockout of Rabaptin5 doubled it. Re-expression of the wild-type protein in knockout cells recovered wild-type levels of survival, while expression of the mutant Rabaptin5-AAA that is unable to bind ATG16L1 did not.

These results suggest that Rabaptin5 recruits autophagy to the damaged membrane of SCVs and contributes to killing of phagocytosed *Salmonella*. To confirm this concept, bacteria infecting the different HEK293A cell lines were analyzed by immunofluorescence microscopy for colocalization with TfR as a marker for early endosomal membrane identity and for LC3B as an autophagy marker (Figure 7C). Right after infection, while virtually all bacteria were intracellular, more than half of them were negative for both markers. This fraction rapidly declined for the next 15 min and the fraction in a TfR-positive environment with or without LC3B increased significantly, consistent with increasing acquisition of endosomal markers of the SCVs by fusion with early endosomes.

In wild-type HEK293A cells and Rabaptin5-overexpressing HEK^+Rbpt5^ cells, the predominant population 5–30 min after infection was that of *Salmonella* in double-positive (TfR+/LC3B+) SCV–autophagosomes (Figure 7C). In Rabaptin5 knockout cells, this population was reduced in favor of a majority of TfR single-positive SCVs as a result of reduced autophagy initiation. Expression of Rabaptin5-AAA did not change this situation, confirming that the interaction with ATG16L1 is required to initiate autophagy of SCVs. At the same time, it shows that the phenotype of the knockout cells is not just the result of reduced endosome maturation in the absence of Rabaptin5, since the AAA mutant is functional for its other endosomal functions. In FIP200 knockout cells TfR+/LC3B+ SCVs were also reduced, as expected.

In summary, Rabaptin5, in addition to its function as a regulator of early endosome identity and endosomal maturation, plays an unexpected role in initiating autophagy of damaged early endosomes and bacteria-containing phagosomes via interaction with FIP200 and ATG16L1.

## DISCUSSION

Previously, Rabaptin5 had been characterized as a regulator of Rab5 activity by complex formation with Rabex5, the GDP/GTP exchange factor of Rab5. In addition it had been shown to bind to Rab4·GTP and Rab5·GTP, the hallmark Rab proteins of early endosomes (Horiuchi et al., 1997; Kälin et al., 2015; 2016; Lippé et al., 2001; Mattera et al., 2006). Rabaptin5 thus was known to contribute to endosome identity and maturation (Huotari and Helenius, 2011). Silencing of Rabaptin5 has no effect on cell viability, most likely because Rabex5 can recruit independently in the absence of Rabaptin5 to endosomes and activate Rab5 (Mattera and Bonifacino, 2008). However, silencing Rabaptin5 or reduced expression of Rabaptin5 due to hypoxia-activated hypoxia-inducible factor (HIF) or overexpressed histone deacetylase 6 (HDAC6) in gastric cancer has been found to delay early endosome fusion, reduce EGF receptor degradation, and thereby prolong receptor signalling (Park et al., 2014; Wang et al., 2009). Consistent with these functions, we found here that the number of lysosomes, the endpoint organelles of the endosomal pathway, was increased by overexpression and reduced upon knockout of Rabaptin5. This phenotype was rescued by re-expression of wild-type as well as of Rabaptin5-AAA that is unable to bind ATG16L1, indicating that the mutant protein performs its endosomal functions normally.

In this study, we identified FIP200 and ATG16L1 as novel interactors of Rabaptin5 and thus an additional unexpected role for Rabaptin5 in autophagy. Endosomes – specifically recycling endosomes – were previously implicated as a source of membranes and of ATG9 and ATG16L1 for growing phagophores (Knævelsrud et al., 2013; Puri et al., 2013; Popovic and Dikic, 2014; Søreng et al., 2018). Rab11A-positive recycling endosomes were found to act as a platform for autophagosome assembly mediated by the interaction of WIPI2 with PI3P and Rab11A (Puri et al., 2018). These recycling endosomes were demonstrated to be distinct from Rab5- and thus also from Rabaptin5-positive early sorting endosomes. While Rab11A was shown to be broadly important for starvation-induced autophagy as well as mitophagy (Puri et al., 2018), silencing of Rabaptin5 did not affect Torin1-induced autophagy or lysophagy, but specifically inhibited autophagy induced by chloroquine or monensin and of SCVs. This specificity argues against a defect in sorting endosomes due to silencing Rabaptin5 to generally inhibit autophagy.

On the contrary, there is evidence that defects in autophagy affect sorting endosome function: cells lacking ATG7 or ATG16L1 were shown to perturb EGF receptor endocytic trafficking and to abrogate EGF receptor signalling (Fraser et al., 2019). Damaged, galectin8-positive early endosomes accumulated in the absence of autophagy, like during chemical disruption with monensin. These findings suggest autophagy of early endosomes to be a housekeeping activity to keep sorting endosomes functional. Rabaptin5 is a good candidate to target autophagy to this compartment.

We found Rabaptin5 to be essential to recruit the autophagy machinery to damaged endosomes by binding to FIP200 and ATG16L1. Rabaptin5 binds to the WD domain of ATG16L1 via an interaction motif ^507^**Y**RLVS**E**TE**W**NL**L**^518^ based on a conserved consensus initially identified in TMEM59, NOD2, and TLR2 to promote autophagy in the context of bacterial phagosomes (Boada-Romero et al., 2013). Rabaptin5 with three of the key residues in this sequence mutated to alanines could not rescue chloroquine-induced WIPI2 and LC3B puncta beyond the level of the Rabaptin5 knockout and did not induce colocalization with WIPI2 or ATG16L1. This suggests that, in the context of damaged early endosomes, Rabaptin5 makes an essential contribution to recruit ATG16L1. The importance of the interaction with FIP200 could not be assessed, since deletion of the interacting segment in Rabaptin5 (residues 547–666; CC2-1) abolishes Rabex5-binding and membrane recruitment (Kälin et al., 2015) and in FIP200 destabilized complex formation with ATG13 and ULK1.

Chloroquine-treatment caused Rabaptin5-positive endosomes to swell and brake, recruiting galectin3, ubiquitination, FIP200, WIPI2, and ATG16L1 within 30 min. At longer time points, also LC3B-positive autophagosomes accumulated. Induction of WIPI2 puncta was strongly reduced by silencing of Rabaptin5, FIP200, ATG13, or by inhibition of ULK1/2. LC3B autophagosomes/autolysosomes were also reduced, but to a lesser extent, most likely because of their accumulation due to inhibition of autophagic flux. The involvement of the ULK1–FIP200 initiator complex and of WIPI2 is in agreement with canonical selective autophagy upon chloroquine treatment, similar to lysophagy induced by LLOMe treatment (Chauhan et al., 2016; Maejima et al., 2013). However, it differs from a noncanonical LAP-like mechanism that was observed for entotic vacuoles or latex bead-containing phagosomes as model organelles (Fletcher et al., 2018; Florey et al., 2015; Jacquin et al., 2017). Upon chloroquine or monensin treatment, LC3 was recruited directly to the single membrane vacuoles without PI3P production or WIPI involvement, without ubiquitination, and independently of ATG13 and of the FIP200-binding domain of ATG16L1. Chloroquine and monensin appear to trigger different responses on large phagosomal structures and entotic vacuoles than on regular early endosomes.

*Salmonella* that are phagocytosed by professional phagocytes like macrophages are the target of LAP (Masud et al., 2019). However, *Salmonella* also has the ability to actively infect other cell types such as epithelial cells and fibroblasts, triggering uptake into SCVs that are the target of canonical autophagy (LaRock et al., 2015).

*Salmonella* manipulates its host by injecting effectors into the host cytosol via two Type III secretion systems (T3SS). T3SS-1 is required for internalization into SCVs. As SCVs are further acidified, T3SS-2 is induced and its effectors mediate maturation to *Salmonella*-induced filaments, where bacteria proliferate. However, T3SS-1 also damages the SCV membrane. This may result in complete release of bacteria into the cytosol where they hyperproliferate and become direct targets of ubiquitination and autophagy. In contrast, damaged SCVs are recognized by selective autophagy in a similar manner as damaged endosomes or lysosomes: exposure of lumenal protein domains and glycans trigger the binding of galectin3 and 8 (Fujita et al., 2013; Thurston et al., 2012), poly-ubiquitination, and p62 and NDP52 recruitment (Thurston et al., 2012; Zheng et al., 2009). SCVs are surrounded by phagophores in a process requiring FIP200 and ATG9L1 (e.g. Fujita et al., 2013; Huang and Brumell, 2014; Kageyama et al., 2011; Kishi-Itakura et al., 2020).

Because of the similarities between autophagy of SCVs and of damaged endosomes, we tested a potential role of Rabaptin5 in the early phase of *Salmonella* infection. We found that autophagic elimination of *Salmonella* during the first hour post infection, when the SCVs most resemble early endosomes, strongly depended on Rabaptin5 and its ability to bind ATG16L1. In wild-type HeLa and HEK293A cells, 70–80% of internalized bacteria were killed within the first hour. Knockdown or knockout of Rabaptin5 reduced this number to approximately 50%, i.e. to the same extent as silencing of FIP200 did. The same effect was observed, when wild-type Rabaptin5 was replaced by a triple mutant defective in its interaction with ATG16L1.

Interestingly, Kageyama et al. (2011)observed that LC3-recruitment to SVCs was not blocked in the absence of the ULK1/FIP200 complex, ATG9L1, or ATG14L, or in the presence of wortmannin, suggesting a noncanonical mechanism of recruiting ATG16L1 to SCVs in the absence of the canonical machinery. This alternative mechanism may be responsible for the residual killing of bacteria we observed in the absence of Rabaptin5 or FIP200.

Our study shows that Rabaptin5 has dual functions: in addition to its role as a regulator of early endosome identity and endosome maturation, it contributes to initiate and promote autophagy, when endosomes are damaged spontaneously, by chemical agents, or by pathogens.

## MATERIALS AND METHODS

### Antibodies and reagents

Mouse anti-transferrin receptor (OKT8 mouse monoclonal hybridoma, kind gift of Dr. H. Farhan, 1:1000), mouse anti-Rbpt5 (610676, BD Transduction Laboratories, 1:1000) and rabbit anti-Rabaptin5 antibodies (NBP1-47285, Novus Biologicals, 1:500) were used for immunofluorescence and Western blotting, rabbit anti-FIP200 (12436S, Cell Signaling, 1:1000) was used for Western blot and co-immunoprecipitation. Mouse anti-tubulin (kind gift of Dr. H. Farhan, 1:1000) was used for immunoblotting. Rabbit anti-Galectin3 (1:1000; Abcam ab31707), rabbit anti-ATG16L1 (8089, Cell Signaling, 1:1000), rabbit anti-MYC (GTX29106, GeneTEX, 1:5000) rabbit anti-mCherry (PA5-34974, Invitrogen, 1:1000) and rabbit EEA1were (1:1000; Abcam Ab2900 used for co-immunoprecipitations. Mouse anti-WIPI2 (ab105459, Abcam, 1:500) and rabbit anti LC3B (3868S, Cell Signaling, 1:1000 WB, 1:400 IF) were used for Western blot and immunofluorescence. Rabbit anti-TRIM16 (NB100-59772, Novus Biologicals, 1:1000), rabbit anti-ULK1 (#8054S, Cell Signaling, 1:1000) and rabbit anti ATG13 (#13468S, Cell Signaling, 1:1000) were used for immunoblotting. Mouse anti-ubiquitin (SC-8017, Santa Cruz, 1:500) and mouse anti-ALIX (634502, BioLegend, 1:400) were used for immunofluorescence. Alexa-Fluor-488- or Alexa-Fluor-568-tagged donkey anti-mouse- or anti-rabbit-immunoglobulin antibodies (A32766, A32790, A10042, A10037, Molecular Probes, 1:200) and Alexa-Fluor-633 goat anti mouse (A-21050, Molecular Probes, 1:200) were used as secondary antibodies for immunofluorescence, and horseradish-peroxidase-coupled goat anti-mouse- or anti-rabbit-immunoglobulin antibodies (A0168, A0545, Sigma Immunochemicals, 1:5000) in combination with the enhanced chemiluminescence reaction kit (Amersham Pharmacia Biotech).

Monensin, MRT68921, and LLOME were from Sigma Aldrich (M5273, SML1644, and L7393, respectively), chloroquine from Serva (16919).

### Plasmids and mutagenesis

Plasmids of human Rabaptin5, RFP-Rab5 (pSI-AAR6_Rab5), and GFP-Rab7 (pSI-AAL6_Rab7) were described in Kälin et al. (2015). The cDNA of human FIP200 was purchased from OriGene (SC114884), pmCherry_Gal3 was a gift from Hemmo Meyer (Addgene plasmid #85662) (Papadopoulos et al., 2017), pSpCas9(BB)-2A-GFP (PX458) was a gift from Feng Zhang (Addgene plasmid #48138) (Ran et al., 2013), and mCherry-ATG16L1 was a kind gift from Sharon Tooze, London. Mutations were introduced by PCR mutagenesis. HeLa cells stably expressing GFP-CHMP4B were a kind gift by Prof. A. Hyman (MPI, Dresden).

### Cell culture and transfection

HeLaα and HEK293A cells were grown in Dulbecco’s modified Eagle’s medium (DMEM) supplemented with 10% fetal calf serum (FCS), 100 units/ml penicillin, 100 units/ml streptomycin, and 2 mM L-glutamine at 37°C at 7.5% CO_2_.

To generate Rabaptin5- and FIP200-knockout cell lines in HEK293A cells, the gRNA sequences AGAGTGTGTACCTACAGTGC (Rabaptin5) and TGGGCGCCTCACCGTCAGGC (FIP200) extended with Bpi I overhangs were cloned into pSpCas9(BB)-2A-GFP, digested with Bpi I (Thermo Scientific, FD1014). 24 h after transfection, the plasmid containing cells were selected by fluorescence-activated cell sorting based on GFP expression. After 10 days of culture, single cells were sorted again for loss of GFP expression and expanded. Gene inactivation was confirmed by immunoblotting

Transient transfections were performed using Fugene (Promega) or JetPRIME (Polyplus) and cells analyzed after 2 days. For stable expression, cells were transfected with the respective pcDNA3 constructs and subsequently selected with 2 mg/ml G418 (Geneticin from Thermo Fisher Scientific; 3 mg/ml for 10 days). Surviving cells were expanded in culture medium containing G418.

For knockdown experiments, siRNAs were purchased from Dharmacon Thermo Fisher Scientific (nontargeting siControl, D-001810-10-05; Rabaptin5 siRABEP1, L-017645-01; FIP200 siRB1CC1, M-021117-01-0005; siATG16L1, J-021033-11-0010; siATG13, L-020765-01; siULK1, L-005049-00; siWIPI2, J-020521-09-0005). Cells were transfected with 20 nM siRNA and HiPerFect (QIAGEN), and used after 3 days.

### Immunofluorescence microscopy and quantitation

Cells on coverslips were fixed with 3% paraformaldehyde (PFA) for 10 min at room temperature, quenched for 5 min with 50 mM NH_4_Cl, washed with PBS, permeabilized with 0.1% Triton X-100 for 10 min, blocked with 3% BSA in PBS for 15 min, incubated for 1 h with primary antibodies in PBS with BSA, washed, and stained for 30 min with fluorescently tagged secondary antibodies in PBS with BSA. Coverslips were mounted in Fluoromount-G (Southern Biotech).

For LC3B and WIPI2 staining, fixation was performed with methanol or, when co-staining for Rabaptin5, with PFA/methanol. In the first case, cells were fixed for 15 min in –20°C methanol, permeabilized for 10 min with 0.3% Triton X-100, blocked for 30 min with 3% BSA in PBS, and incubated overnight with the primary antibody at 4°C. In the second case, cells were first fixed in 3% PFA for 1 min and then for 15 min in 100% – 20°C methanol, permeabilized for 10 min with 0.1% Triton X-100, blocked for 15 min with 3% BSA in PBS, and incubated overnight with the primary antibody at 4°C. Secondary antibody was incubated in 3% BSA for 30 min at room temperature. PFA/methanol fixation was also used to co-stain Rabaptin5 and ALIX in HEK293A cells, HEK293A-FIP200-KO cells and GFP-CHMP4B Hela cells.

Images were acquired using an LSM700 confocal microscope or an API Delta Vision Core microscope with a 63x lens. All experiments were repeated at least three times. Images were analyzed with Fiji, using the JACoP plugin to determine Manders’ colocalization coefficients. Quantitations were based on images of 11 fields typically containing 3–8 cells. To quantify colocalization between Rabaptin5 and mCherry-ATG16L1, transfected cells were fixed with PFA/methanol as above and stained for Rabaptin5. Images of at least 20 mCherry-ATG16L-expressing cells were analyzed per experiment.

To estimate the number of lysosomes, cells were loaded overnight with 450 nM Lysotracker Green (from Molecular Probes), fixed in 3% PFA, and imaged. Images for at least 15 cells per sample were quantified using Fiji.

To evaluate autophagic flux, HEK293A and Rabaptin5-KO cells were transduced with the Premo Autophagy Tandem Sensor RFP-GFP-LC3B Kit (Thermo Fisher Scientific) for 48 h and incubated with or without HBSS starvtion media or with 250 nM Torin1 for 3 h before fixation. GFP/RFP double-positive autophagosomes and GFP-negative/RFP-positive autolysosomes were quantified by fluorescence microscopy.

### Co-immunoprecipitation

Cells were lysed with lysis buffer (0.5% Na-deoxycholate, 1% Triton X-100, 2 mM PMSF, protease inhibitor cocktail). Post-nuclear supernatants were incubated with antibody overnight at 4°C. Antigen–antibody complexes were collected with protein A-Sepharose for 2 h at 4°C, washed three times with lysis buffer, and subjected to immunoblot analyses.

### Salmonella infection and immunofluorescence analysis

Wild-type *Salmonella enterica* serovar Typhimurium strain SL1344 was cultured in Luria broth (LB) overnight at 37°C with shaking, followed by dilution into 10 ml of fresh LB (1:33), and continued to grow under the same conditions for 3 h. One milliliter of bacteria was then centrifuged at 8,000×g for 2 min and resuspended in 1 ml of PBS. This suspension was diluted into DMEM + 10% FCS (no antibiotics) and added directly to HeLa or HEK293A cell lines to a multiplicity of infection of 100. The infected cell monolayers were centrifuged at 500×g for 5 min at 37°C to synchronize the infection and incubated 10 min at 37°C. The monolayers were then washed three times in PBS and then incubated in fresh culture medium with 100 µg/ml gentamycin. At 1 h post infection, the gentamycin concentration was reduced to 10 µg/ml. At designated time points, cells were washed three times with PBS and lysed in 500 µl of lysis buffer (PBS with 1% Triton X-100). Serial dilutions were plated onto LB agar plates (with 90 µg/ml streptomycin) to determine bacterial live cell count.

For microscopy, cells were infected with GFP-expressing *Salmonella* SL1344 as above, fixed after different times by the methanol fixation protocol, and incubated with anti-TfR and anti-LC3B antibodies in PBS with 3% BSA overnight at 4°C, followed by fluorescent secondary antibodies for 30 min at room temperature. Images were acquired using a Zeiss LSM700 confocal microscope and analysed in Fiji. To determine the infection rate, HeLa cells infected with GFP-expressing *Salmonella* SL1344 as above were fixed in methanol and stained for TfR and LC3B. Z-stacks if images were acquired on an API Delta Vision Core microscope using the 20x lens and a spacing of 0.13 μm. A range of 5’000 – 18’000 cells/sample were imaged on 5-7 different areas on the coverslip, deconvoluted and stitched together on a SoftWoRx Imaging Workstastion. The final 3D pictures were analyzed with Fiji.

### Yeast two-hybrid screen and interaction domain identification

The pB29-Rbpt5(1–862)-LexA bait plasmid was used to screen a random-primed HeLa cells_RP1 cDNA library cloned into the pP6-Gal4-AD plasmid using a high-throughput proprietary yeast two-hybrid-based technology (ULTImate Y2H Screen; Hybrigenics, Paris, France). To identify the FIP200 interaction segment in Rabaptin5, interaction domain mapping analysis of different fragments in pB29-Rbpt5(#)-LexA was performed with pP6-Gal4AD-FIP200(257–444) (Hybrigenics).

## ACKNOWLEDGMENTS

We thank Drs. Mauricio Rosas Ballina and Dirk Bumann (Biozentrum) for materials and assistance with the *Salmonella* experiments, Drs. Kai Schleicher, Alexia Loynton-Ferrand and the Biozentrum Imaging Core Facility, and Janine Bögli of the FACS Core Facility for their support. This work was supported by Grant 31003A-182519 from the Swiss National Science Foundation.

## AUTHOR CONTRIBUTIONS

VM, SS, SK planned and performed experiments and analyzed data. MS planned experiments and analyzed data. VM and MS wrote the manuscript.

## CONFLICT OF INTEREST

The authors declare that they have no conflict of interest.

## SUPPLEMENTARY INFORMATION

**Suppl. Table S1.** Rabaptin5 interactors identified by yeast two-hybrid screen.

| Interaction quality* | Gene name | Protein | Interacting fragment |
| --- | --- | --- | --- |
| A | RABEP1 | Rabaptin5 | 579–691 |
| **A** | **RB1CC1** | **FIP200** | **281–439** |
| A | VIM | Vimentin | 15–144 |
| B | RABGEF1 | Rabex5 | 281–439 |
| B | BARD1 | BRCA1-associated ring domain protein 1 | 47–284 |
| B | GOLGA4 | Golgin A4 | 1116–1281 |
| B | ITSN1 | Intersectin 1 | 554–667 |
| C | SMARCC1 | SWI/SNF complex subunit SMARCC1 | 899–1005 |

\*A–C indicate the confidence level of the interaction from very high to medium.
FIP200, FAK family-interacting protein of 200 kDa; RB1CC1, RB1-inducible coiled-coil protein 1.

**Suppl. Figure S1.**
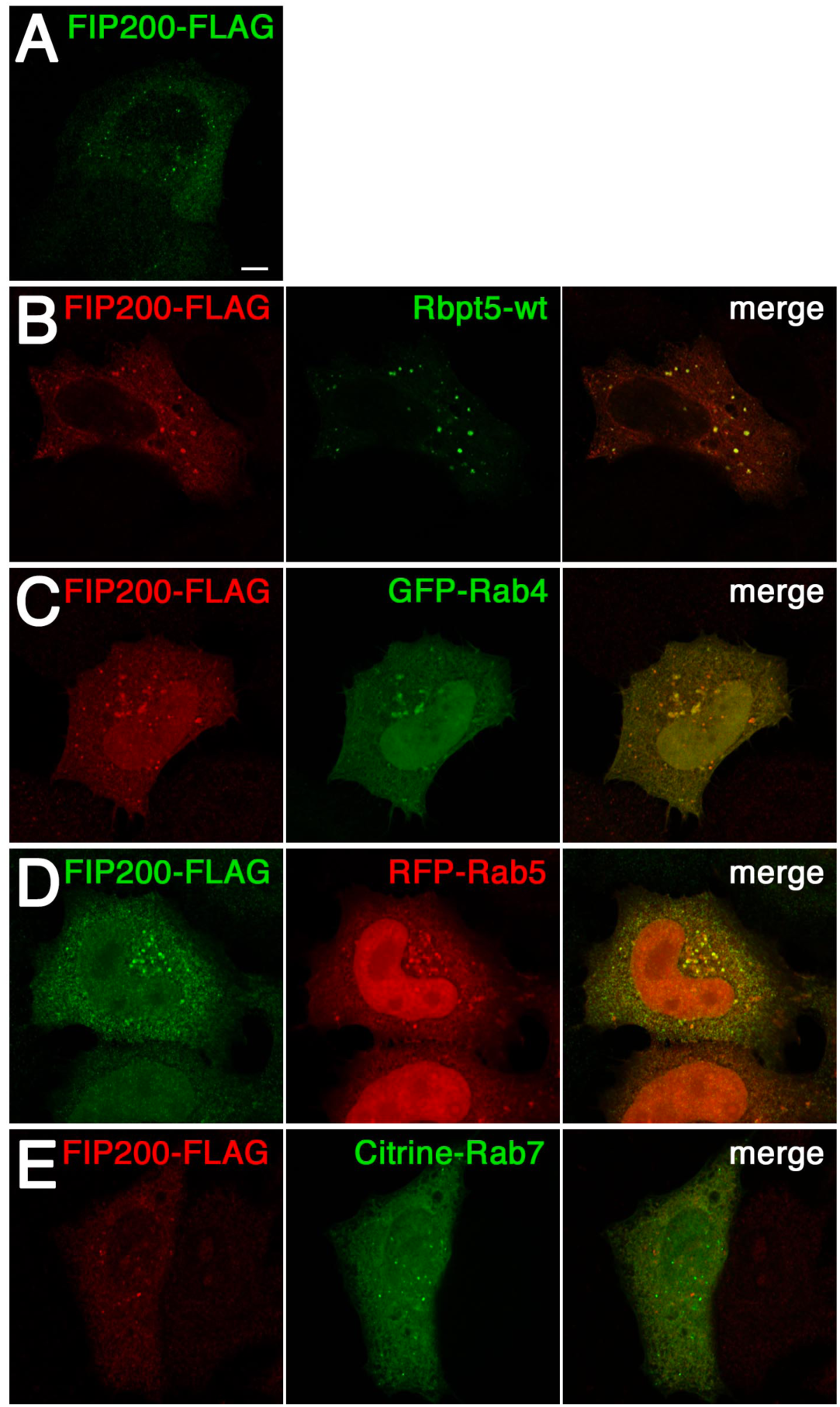
FIP200 colocalizes with endosomal markers. HeLa cells were transfected with FLAG-tagged FIP200 alone (A) or together with Rabaptin5 (Rbpt5-wt) (B), GFP-Rab4 (C), RFP-Rab5 (D), or Citrine-Rab7 (E), fixed after 24 h, and subjected to immunofluorescence microscopy. Bar, 10 µm.

**Suppl. Figure S2.**
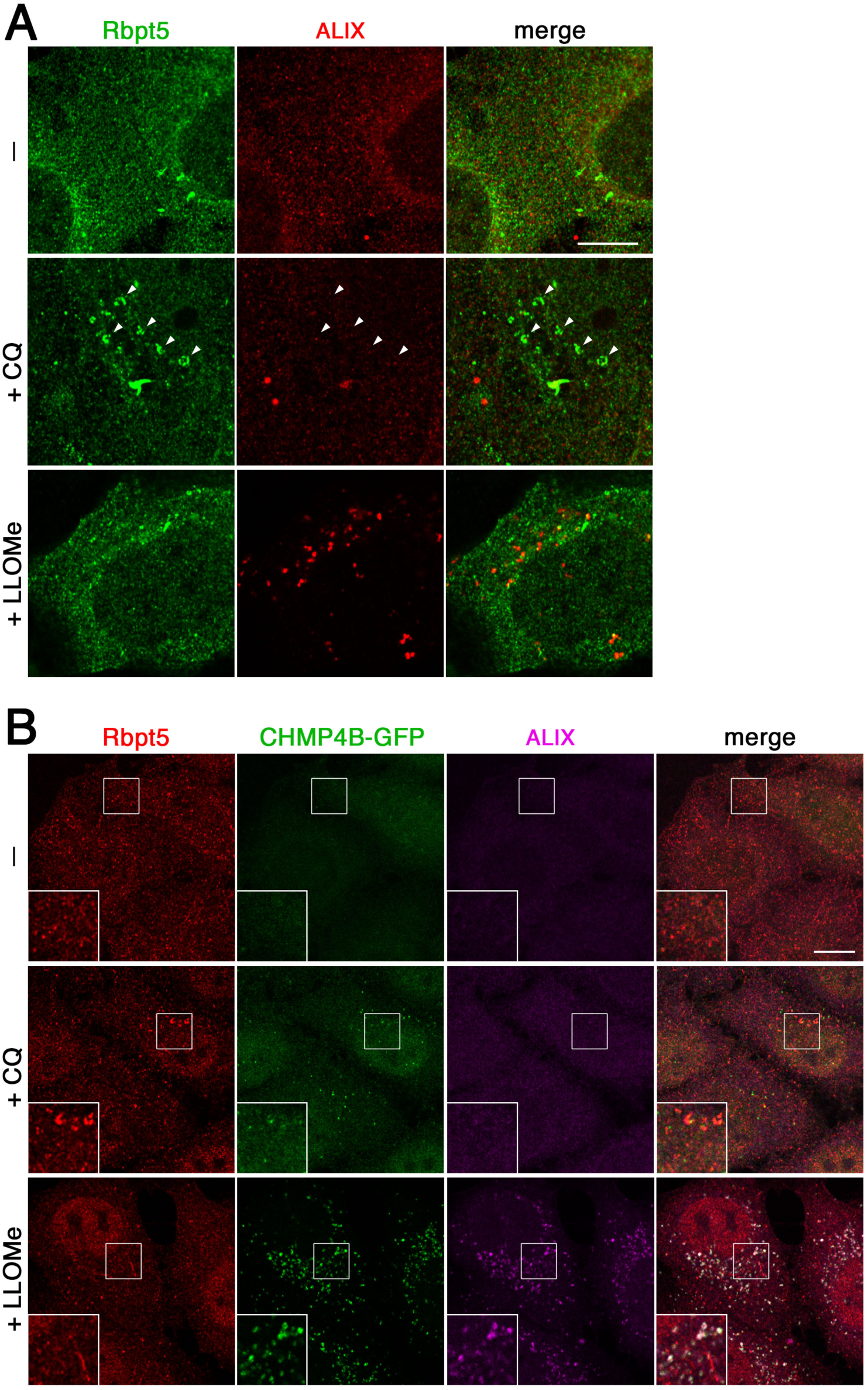
Chloroquine treatment does not recruit ESCRT components to Rabaptin5-positive endosomes. **A**: HEK^+Rbpt5^ cells were treated without (–) or with 60 µM chloroquine (+CQ) or 1 mM LLOMe (+LLOMe) for 30 min and analyzed by immunofluorescence microscopy for Rabaptin5 and ALIX. **B**: HeLa cells stably expressing CHMP4B-GFP were treated as in panel A and analyzed for Rabaptin5, CHMP4B-GFP, and ALIX.

**Suppl. Figure S3.**
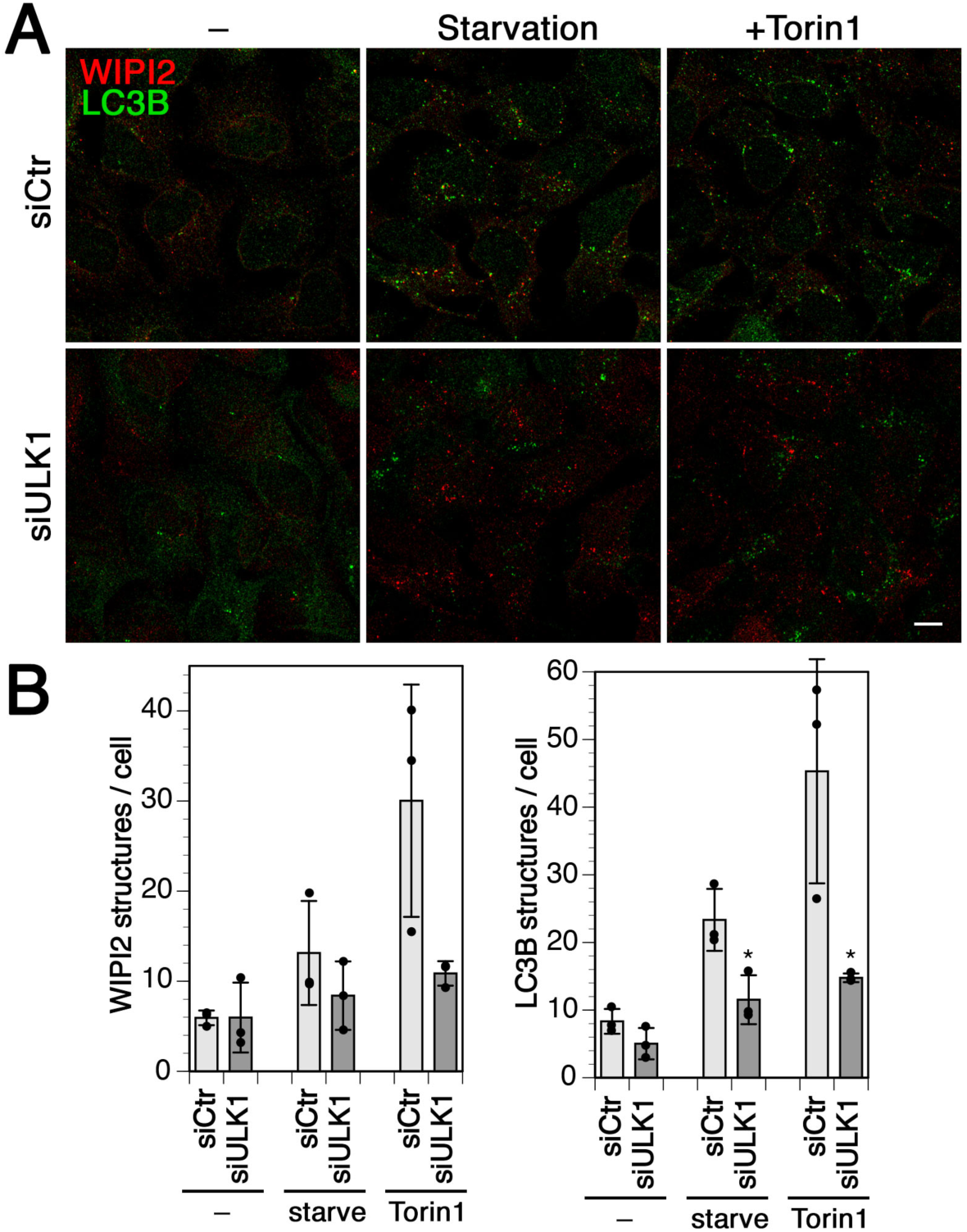
ULK1 knockdown inhibits starvation- and Torin1-induced autophagy. **A:** HEK293A cells were transfected with nontargeting siRNA (siCtr) or siRNAs silencing ULK1 for 72 h, and treated without (–) or with 250 nM Torin1 for 150 min, or starved in HBSS for 150 min. Cells were fixed and immunostained for endogenous WIPI2 and LC3B. Bar, 10 µm. **B and C**: WIPI2 (B) of LC3B puncta per cell (C) were quantified for each condition (mean and standard deviation of three independent experiments; two-tailed Student’s t test: *p < 0.05).

**Suppl. Figure S4.**
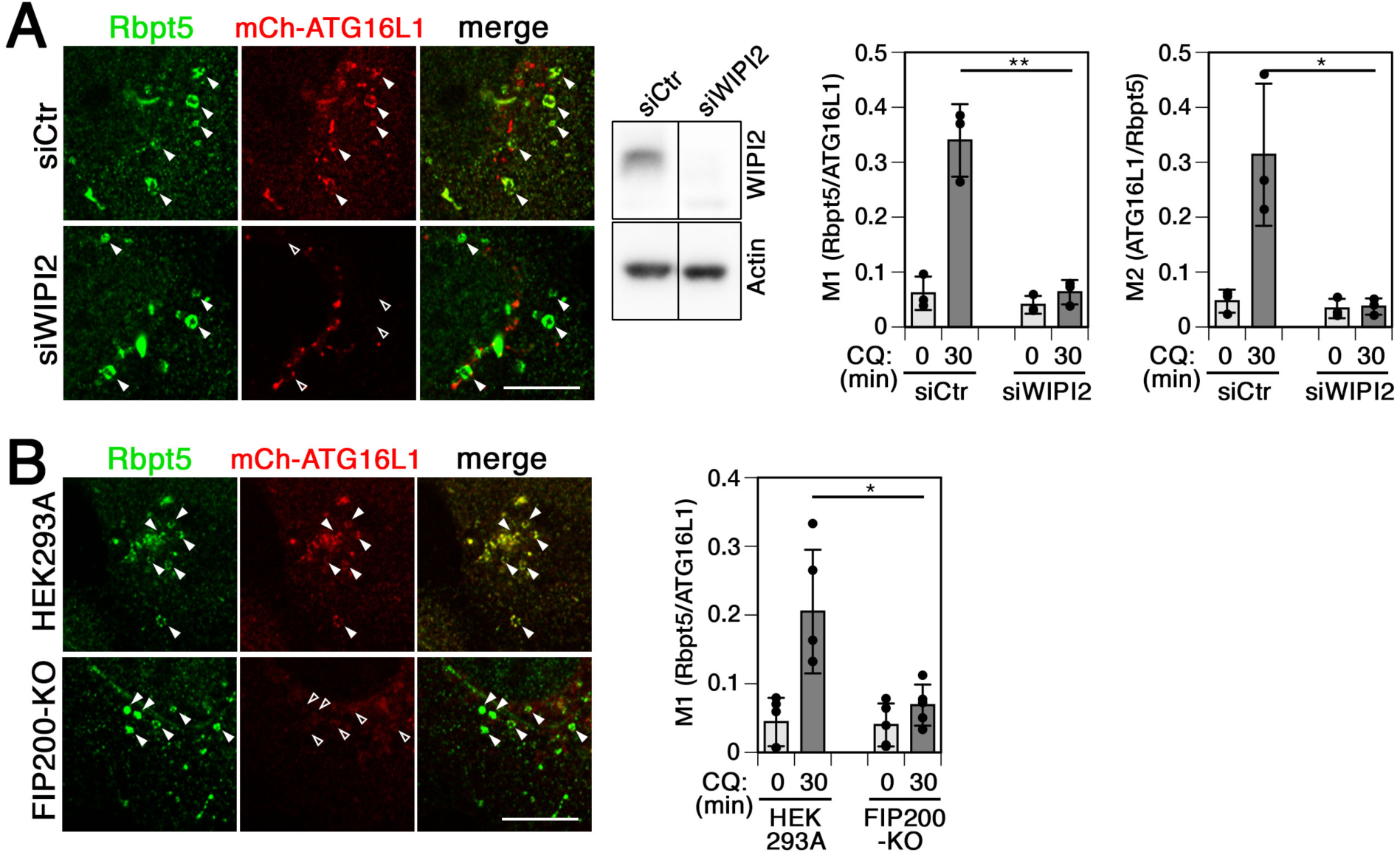
ATG16L1 recruitment to chloroquine-damaged endosomes requires WIPI2 and FIP200. **A**: HEK^+Rbpt5^ cells were transfected with nontargeting siRNA (siCtr) or siRNAs silencing WIPI2 (siWIPI2) for 72 h and with mCherry-ATG16L1 for 24 h. Cells were treated without or with 60 µM chloroquine (+CQ) for 30 min and stained for Rabaptin5 and mCherry-ATG16L1. Fluorescence micrographs of chloroquine-treated cells are shown (left panel). Bar, 10 µm. Arrowheads point out chloroquine-induced enlarged early endosomes. The efficiency of WIPI2 knockdown was assayed by immunoblotting using actin as a loading control (middle panel). Manders’ colocalization coefficients were determined (right panel), M1 showing the fraction of Rabaptin5-positive structures also positive for mCherry-ATG16L1 and M2 showing the inverse. Mean and standard deviation of three independent experiments; ANOVA: *p < 0.05, **p < 0.01. **B**: Parental HEK293A and FIP200 knockout cells (FIP200-KO) were transfected with mCherry-ATG16L1 for 24 h and incubated without or with 60 µM chloroquine (+CQ) for 30 min and stained for Rabaptin5 and mCherry-ATG16L1. Fluorescence micrographs of chloroquine-treated cells are shown (left panel). Bar, 10 µm. Arrowheads point out chloroquine-induced enlarged early endosomes. Manders’ colocalization coefficients were determined (right panel). Mean and standard deviation of five independent experiments; ANOVA: *p < 0.05.

**Suppl. Figure S5.**
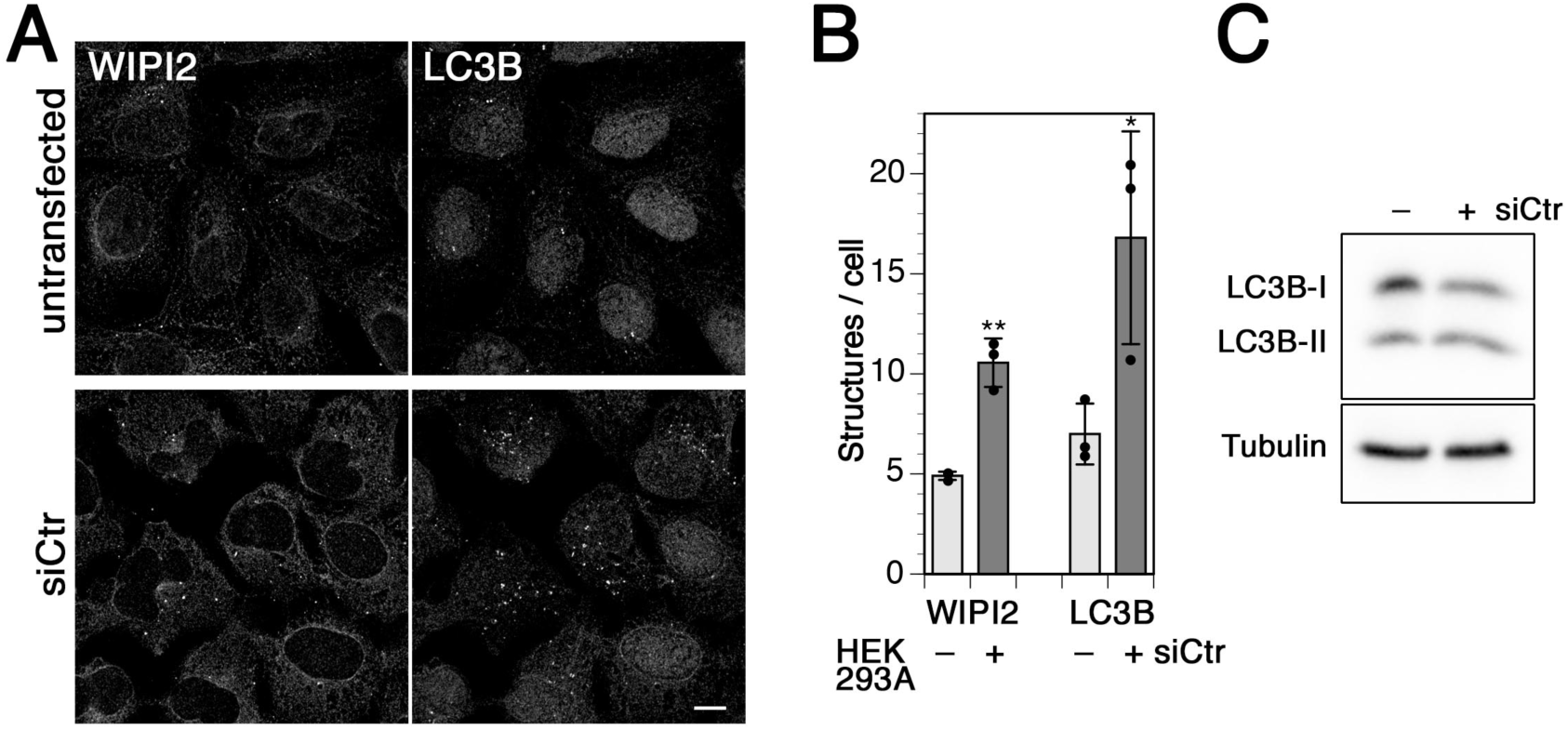
siRNA transfection increases basal autophagy. **A:** HEK293A cells were transfected with nontargeting siRNA (siCtr) for 72 h or not treated. Cells were fixed and immunostained for endogenous WIPI2 and LC3B. Bar, 10 µm. **B:** WIPI2 and LC3B puncta per cell were quantified (mean and standard deviation of three independent experiments; two-tailed Student’s t test: *p < 0.05, **p < 0.01). **C:** Lipidated and non-lipidated LC3B (I and II, resp.) from lysates of HEK293A cells transfected or not with nontargeting siCtr were assayed by immunoblot anlysis.

**Suppl. Figure S6.**
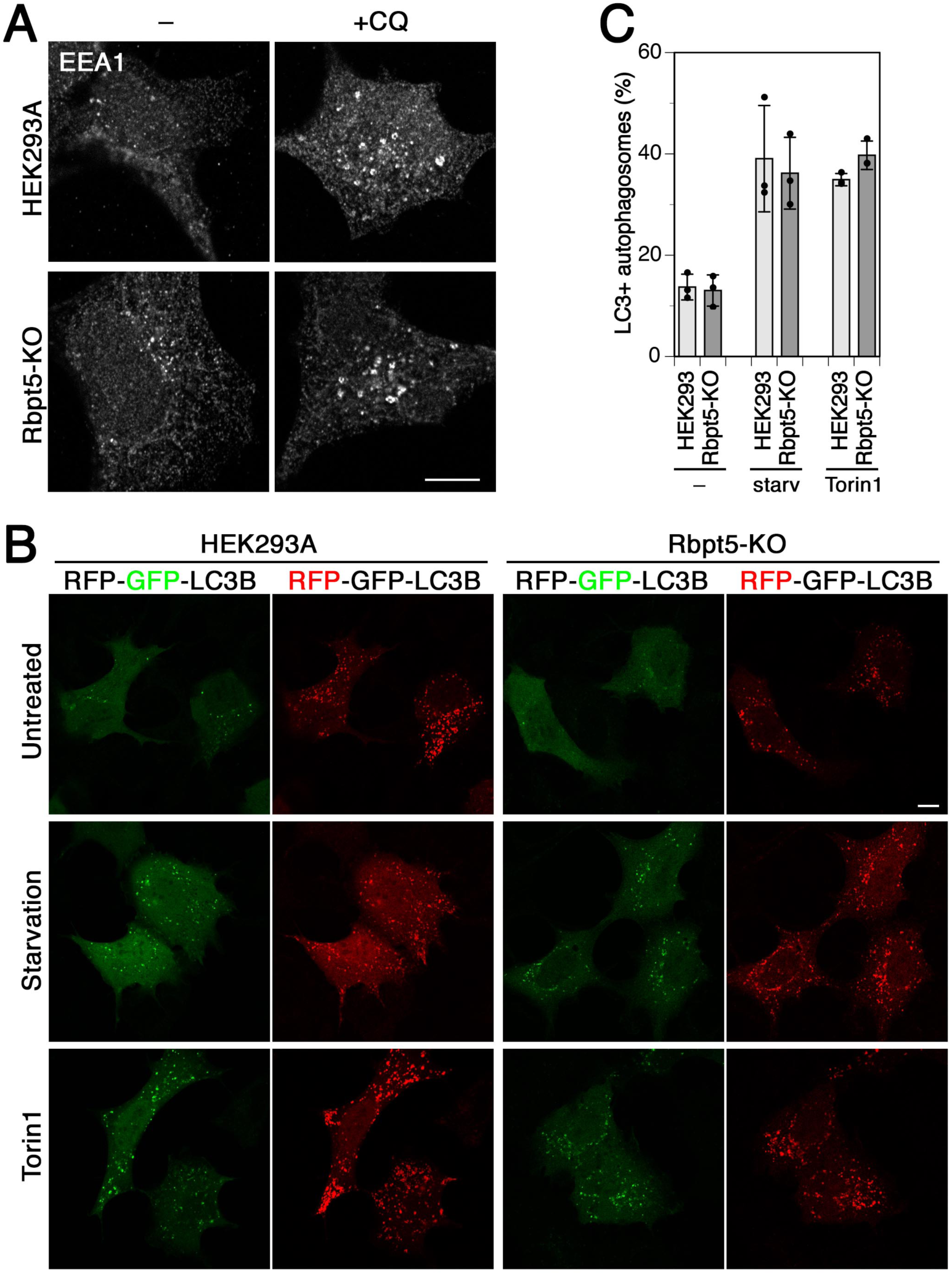
Rabaptin5 knockout does not affect sensitivity of early endosomes to chloroquine or autophagic flux. **A:** Treatment of HEK293A and Rabaptin5-KO cells with 60 µM chloroquine for 30 min similarly leads to swelling of early endosomes positive for EEA1 as detected by immunofluorescence. **B:** HEK293A cells and Rabaptin5-KO cells were transduced to express tandem fluorescent RFP-GFP-LC3, starved in HBSS or treated with 250 nM Torin1 for 150 min, fixed and subjected to fluorescnet microscopy. Images for at least 16 cells per sample were quantified. **C:** The number of autophagosomes (GFP+/RFP+) and autolysosomes (GFP /RFP+) LC3 puncta were counted and autophagosomes plotted as a percentage of the total (mean and standard deviation of three independent experiments).

**Suppl. Figure S7.**
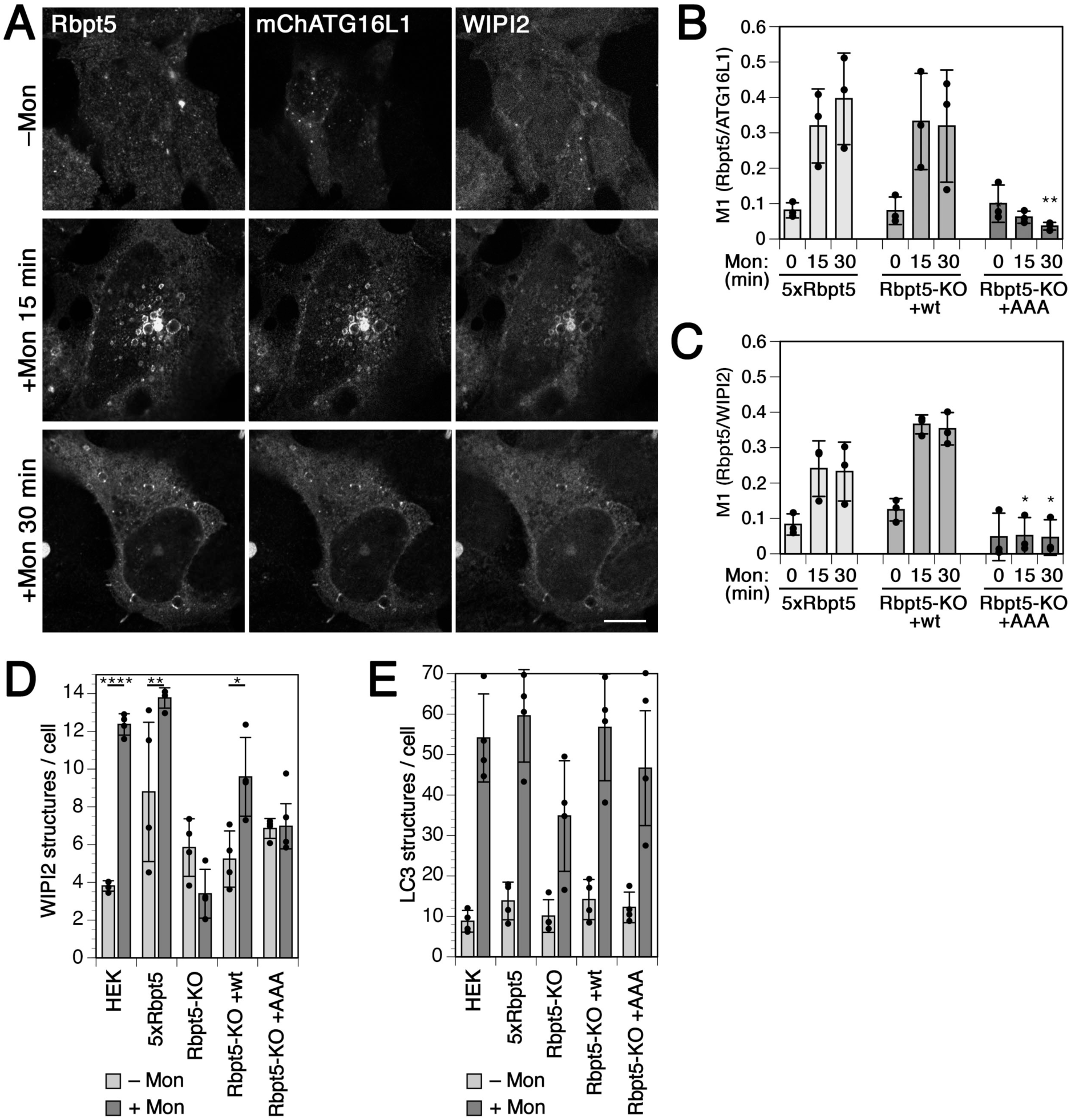
Monensin induces endosomal autophagy dependent on Rabaptin5 and ATG16L1 binding just like chloroquine. **A**: HEK^+Rbpt5^ cells, 24 h after transfection with mCherry-ATG16L1, were treated with 100 µM monensin for 0, 15 and 30 min and stained for Rabaptin5,WIPI2, and mCherry-ATG16L1 to assess their colocalization on swollen early endosomes. Bar, 10 µm. **B** and **C**: Manders’ colocalization coefficients were determined, showing the fraction of Rabaptin5-positive structures also positive for mCherry-ATG16L1 (B) or for WIPI2 (C) (mean and standard deviation of three independent experiments, quantifying ∼40 cells for each sample; ANOVA for Rbpt5-KO+AAA vs. HEK^+Rbpt5^ cells: *p < 0.05, **p < 0.01). **D** and **E**: Wild-type HEK293A cells, HEK^+Rbpt5^ cells, and Rabaptin5-knockout cells without (Rbpt5-KO) or with stable re-expression of wild-type (Rbpt5-KO+wt) or AAA-mutant Rabaptin5 (Rbpt5-KO+AAA) were treated without (–Mon) or with 100 µM monensin for 150 min (+Mon), and analyzed by immunofluorescence microscopy for WIPI2 or LC3B. WIPI2 (D) of LC3B (E) puncta per cell were quantified for each condition (mean and standard deviation of four independent experiments; ANOVA: *p < 0.05, **p < 0.01, ****p < 0.0001).

**Suppl. Figure S8.**
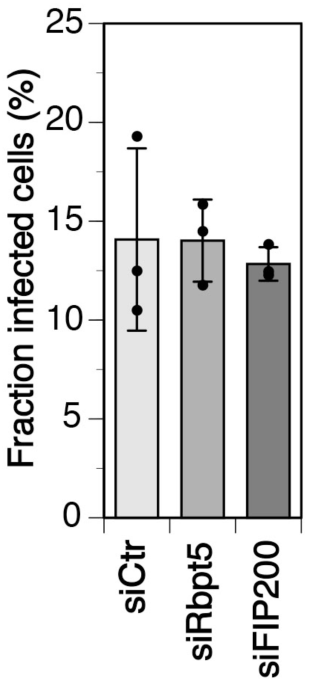
Infection efficiency of Salmonella is not affected upon silencing of Rabaptin5 or FIP200. HeLa cells were transfected with nontargeting siRNA (siCtr) or siRNAs silencing Rabaptin5 (siRbpt5) or FIP200 (siFIP200) for 72 h, infected with *Salmonella* expressing GFP and washed as in Figure 8A and immediately fixed for fluorescence microscopy and stained with anti-transferrin receptor and anti-LC3B. Z-stacks for >5’000cells/sample were acquired and analyzed in Fiji to determine the fraction of infected cells (mean and standard deviation of three independent experiments). The average number of bacteria per infected cell was identical (2.16, 2.11, and 2.12 bacteria per cell transfected with siCtr, siRbpt5, and siFIP200, resp.).

## Notes

### Competing Interest Statement

The authors have declared no competing interest.

### Summary of Updates

A number of additional control experiments were included and the text revised accordingly.

## REFERENCES

Anding, A.L., and Baehrecke, E.H. (2017). Cleaning House: Selective Autophagy of Organelles. Dev Cell 41, 10–22.

Bakula, D., Müller, A.J., Zuleger, T., Takacs, Z., Franz-Wachtel, M., Thost, A.-K., Brigger, D., Tschan, M.P., Frickey, T., Robenek, H., et al. (2017). WIPI3 and WIPI4 β-propellers are scaffolds for LKB1-AMPK-TSC signalling circuits in the control of autophagy. Nat Commun 8, 15637.

Bento, C.F., Renna, M., Ghislat, G., Puri, C., Ashkenazi, A., Vicinanza, M., Menzies, F.M., and Rubinsztein, D.C. (2016). Mammalian Autophagy: How Does It Work? Annu Rev Biochem 85, 685–713.

Birgisdottir, Å.B., and Johansen, T. (2020). Autophagy and endocytosis - interconnections and interdependencies. J Cell Sci 133, jcs228114.

Boada-Romero, E., Letek, M., Fleischer, A., Pallauf, K., Ramón-Barros, C., and Pimentel-Muiños, F.X. (2013). TMEM59 defines a novel ATG16L1-binding motif that promotes local activation of LC3. EMBO J 32, 566–582.

Chaikuad, A., Koschade, S.E., Stolz, A., Zivkovic, K., Pohl, C., Shaid, S., Ren, H., Lambert, L.J., Cosford, N.D.P., Brandts, C.H., et al. (2019). Conservation of structure, function and inhibitor binding in UNC-51-like kinase 1 and 2 (ULK1/2). Biochem J 476, 875–887.

Chauhan, S., Kumar, S., Jain, A., Ponpuak, M., Mudd, M.H., Kimura, T., Choi, S.W., Peters, R., Mandell, M., Bruun, J.-A., et al. (2016). TRIMs and Galectins Globally Cooperate and TRIM16 and Galectin-3 Co-direct Autophagy in Endomembrane Damage Homeostasis. Dev Cell 39, 13–27.

Dikic, I., and Elazar, Z. (2018). Mechanism and medical implications of mammalian autophagy. Nat Rev Mol Cell Biol 19, 349–364.

Dooley, H.C., Razi, M., Polson, H.E.J., Girardin, S.E., Wilson, M.I., and Tooze, S.A. (2014). WIPI2 links LC3 conjugation with PI3P, autophagosome formation, and pathogen clearance by recruiting Atg12-5-16L1. Mol Cell 55, 238–252.

Dudley, L.J., Cabodevilla, A.G., Makar, A.N., Sztacho, M., Michelberger, T., Marsh, J.A., Houston, D.R., Martens, S., Jiang, X., and Gammoh, N. (2019). Intrinsic lipid binding activity of ATG16L1 supports efficient membrane anchoring and autophagy. EMBO J 38, e100554.

Fletcher, K., Ulferts, R., Jacquin, E., Veith, T., Gammoh, N., Arasteh, J.M., Mayer, U., Carding, S.R., Wileman, T., Beale, R., et al. (2018). The WD40 domain of ATG16L1 is required for its non-canonical role in lipidation of LC3 at single membranes. EMBO J 37, e97840.

Florey, O., Gammoh, N., Kim, S.E., Jiang, X., and Overholtzer, M. (2015). V-ATPase and osmotic imbalances activate endolysosomal LC3 lipidation. Autophagy 11, 88–99.

Fraser, J., Simpson, J., Fontana, R., Kishi-Itakura, C., Ktistakis, N.T., and Gammoh, N. (2019). Targeting of early endosomes by autophagy facilitates EGFR recycling and signalling. EMBO Rep 20, e47734.

Fujita, N., Morita, E., Itoh, T., Tanaka, A., Nakaoka, M., Osada, Y., Umemoto, T., Saitoh, T., Nakatogawa, H., Kobayashi, S., et al. (2013). Recruitment of the autophagic machinery to endosomes during infection is mediated by ubiquitin. J Cell Biol 203, 115–128.

Gammoh, N. (2020). The multifaceted functions of ATG16L1 in autophagy and related processes. J Cell Sci 133, jcs249227.

Hara, T., Takamura, A., Kishi, C., Iemura, S.-I., Natsume, T., Guan, J.-L., and Mizushima, N. (2008). FIP200, a ULK-interacting protein, is required for autophagosome formation in mammalian cells. J Cell Biol 181, 497– 510.

Heckmann, B.L., Boada-Romero, E., Cunha, L.D., Magne, J., and Green, D.R. (2017). LC3-Associated Phagocytosis and Inflammation. J. Mol. Biol. 429, 3561–3576.

Horiuchi, H., Lippé, R., McBride, H.M., Rubino, M., Woodman, P., Stenmark, H., Rybin, V., Wilm, M., Ashman, K., Mann, M., et al. (1997). A novel Rab5 GDP/GTP exchange factor complexed to Rabaptin-5 links nucleotide exchange to effector recruitment and function. Cell 90, 1149–1159.

Huang, J., and Brumell, J.H. (2014). Bacteria-autophagy interplay: a battle for survival. Nat Rev Microbiol 12, 101–114.

Huotari, J., and Helenius, A. (2011). Endosome maturation. EMBO J 30, 3481–3500.

Jacquin, E., Leclerc-Mercier, S., Judon, C., Blanchard, E., Fraitag, S., and Florey, O. (2017). Pharmacological modulators of autophagy activate a parallel noncanonical pathway driving unconventional LC3 lipidation. Autophagy 13, 854–867.

Jia, J., Abudu, Y.P., Claude-Taupin, A., Gu, Y., Kumar, S., Choi, S.W., Peters, R., Mudd, M.H., Allers, L., Salemi, M., et al. (2018). Galectins Control mTOR in Response to Endomembrane Damage. Mol Cell 70, 120– 135.e128.

Jia, J., Claude-Taupin, A., Gu, Y., Choi, S.W., Peters, R., Bissa, B., Mudd, M.H., Allers, L., Pallikkuth, S., Lidke, K.A., et al. (2020). Galectin-3 Coordinates a Cellular System for Lysosomal Repair and Removal. Dev Cell 52, 69–87.e8.

Kageyama, S., Omori, H., Saitoh, T., Sone, T., Guan, J.-L., Akira, S., Imamoto, F., Noda, T., and Yoshimori, T. (2011). The LC3 recruitment mechanism is separate from Atg9L1-dependent membrane formation in the autophagic response against Salmonella. Mol Biol Cell 22, 2290–2300.

Kälin, S., Buser, D.P., and Spiess, M. (2016). A fresh look at the function of Rabaptin5 on endosomes. Small GTPases 7, 34–37.

Kälin, S., Hirschmann, D.T., Buser, D.P., and Spiess, M. (2015). Rabaptin5 is recruited to endosomes by Rab4 and Rabex5 to regulate endosome maturation. J Cell Sci 128, 4126–4137.

Kimura, S., Noda, T., and Yoshimori, T. (2007). Dissection of the autophagosome maturation process by a novel reporter protein, tandem fluorescent-tagged LC3. Autophagy 3, 452–460.

Kirkin, V., and Rogov, V.V. (2019). A Diversity of Selective Autophagy Receptors Determines the Specificity of the Autophagy Pathway. Mol Cell 76, 268–285.

Kishi-Itakura, C., Ktistakis, N.T., and Buss, F. (2020). Ultrastructural insights into pathogen clearance by autophagy. Traffic 21, 310–323.

LaRock, D.L., Chaudhary, A., and Miller, S.I. (2015). Salmonellae interactions with host processes. Nat Rev Microbiol 13, 191–205.

Leidal, A.M., Levine, B., and Debnath, J. (2018). Autophagy and the cell biology of age-related disease. Nat Cell Biol 20, 1338–1348.

Levin, R., Grinstein, S., and Canton, J. (2016). The life cycle of phagosomes: formation, maturation, and resolution. Immunol. Rev. 273, 156–179.

Lippé, R., Miaczynska, M., Rybin, V., Runge, A., and Zerial, M. (2001). Functional synergy between Rab5 effector Rabaptin-5 and exchange factor Rabex-5 when physically associated in a complex. Mol Biol Cell 12, 2219–2228.

Maejima, I., Takahashi, A., Omori, H., Kimura, T., Takabatake, Y., Saitoh, T., Yamamoto, A., Hamasaki, M., Noda, T., Isaka, Y., et al. (2013). Autophagy sequesters damaged lysosomes to control lysosomal biogenesis and kidney injury. EMBO J 32, 2336–2347.

Martinez, J., Almendinger, J., Oberst, A., Ness, R., Dillon, C.P., Fitzgerald, P., Hengartner, M.O., and Green, D.R. (2011). Microtubule-associated protein 1 light chain 3 alpha (LC3)-associated phagocytosis is required for the efficient clearance of dead cells. Proc. Natl. Acad. Sci. U.S.A. 108, 17396–17401.

Martinez, J., Malireddi, R.K.S., Lu, Q., Cunha, L.D., Pelletier, S., Gingras, S., Orchard, R., Guan, J.-L., Tan, H., Peng, J., et al. (2015). Molecular characterization of LC3-associated phagocytosis reveals distinct roles for Rubicon, NOX2 and autophagy proteins. Nat Cell Biol 17, 893–906.

Masud, S., Prajsnar, T.K., Torraca, V., Lamers, G.E.M., Benning, M., Van Der Vaart, M., and Meijer, A.H. (2019). Macrophages target Salmonella by Lc3-associated phagocytosis in a systemic infection model. Autophagy 15, 796–812.

Mattera, R., and Bonifacino, J.S. (2008). Ubiquitin binding and conjugation regulate the recruitment of Rabex-5 to early endosomes. EMBO J 27, 2484–2494.

Mattera, R., Tsai, Y.C., Weissman, A.M., and Bonifacino, J.S. (2006). The Rab5 guanine nucleotide exchange factor Rabex-5 binds ubiquitin (Ub) and functions as a Ub ligase through an atypical Ub-interacting motif and a zinc finger domain. J Biol Chem 281, 6874–6883.

Mauthe, M., Orhon, I., Rocchi, C., Zhou, X., Luhr, M., Hijlkema, K.-J., Coppes, R.P., Engedal, N., Mari, M., and Reggiori, F. (2018). Chloroquine inhibits autophagic flux by decreasing autophagosome-lysosome fusion. Autophagy 14, 1435–1455.

Mercer, T.J., Gubas, A., and Tooze, S.A. (2018). A Molecular Perspective of Mammalian Autophagosome Biogenesis. J Biol Chem 293, 5386–5395.

Morishita, H., and Mizushima, N. (2019). Diverse Cellular Roles of Autophagy. Annu Rev Cell Dev Biol 35, 453–475.

Naslavsky, N., and Caplan, S. (2018). The enigmatic endosome - sorting the ins and outs of endocytic trafficking. J Cell Sci 131, jcs216499.

Papadopoulos, C., Kirchner, P., Bug, M., Grum, D., Koerver, L., Schulze, N., Poehler, R., Dressler, A., Fengler, S., Arhzaouy, K., et al. (2017). VCP/p97 cooperates with YOD1, UBXD1 and PLAA to drive clearance of ruptured lysosomes by autophagy. EMBO J 36, 135–150.

Park, S.J., Kim, J.K., Bae, H.J., Eun, J.W., Shen, Q., Kim, H.S., Shin, W.C., Yang, H.D., Lee, E.K., You, J.S., et al. (2014). HDAC6 sustains growth stimulation by prolonging the activation of EGF receptor through the inhibition of rabaptin-5-mediated early endosome fusion in gastric cancer. Cancer Lett 354, 97–106.

Petherick, K.J., Conway, O.J.L., Mpamhanga, C., Osborne, S.A., Kamal, A., Saxty, B., and Ganley, I.G. (2015). Pharmacological inhibition of ULK1 kinase blocks mammalian target of rapamycin (mTOR)-dependent autophagy. J Biol Chem 290, 11376–11383.

Poteryaev, D., Datta, S., Ackema, K., Zerial, M., and Spang, A. (2010). Identification of the switch in early-to-late endosome transition. Cell 141, 497–508.

Puri, C., Vicinanza, M., Ashkenazi, A., Gratian, M.J., Zhang, Q., Bento, C.F., Renna, M., Menzies, F.M., and Rubinsztein, D.C. (2018). The RAB11A-Positive Compartment Is a Primary Platform for Autophagosome Assembly Mediated by WIPI2 Recognition of PI3P-RAB11A. Dev Cell 45, 114–131.e118.

Radulovic, M., Schink, K.O., Wenzel, E.M., Nähse, V., Bongiovanni, A., Lafont, F., and Stenmark, H. (2018). ESCRT-mediated lysosome repair precedes lysophagy and promotes cell survival. EMBO J 37, e99753.

Ran, F.A., Hsu, P.D., Wright, J., Agarwala, V., Scott, D.A., and Zhang, F. (2013). Genome engineering using the CRISPR-Cas9 system. Nat Protoc 8, 2281–2308.

Randow, F., and Youle, R.J. (2014). Self and nonself: how autophagy targets mitochondria and bacteria. Cell Host & Microbe 15, 403–411.

Shi, X., Yokom, A.L., Wang, C., Young, L.N., Youle, R.J., and Hurley, J.H. (2020). ULK complex organization in autophagy by a C-shaped FIP200 N-terminal domain dimer. J Cell Biol 219, e201911047.

Skowyra, M.L., Schlesinger, P.H., Naismith, T.V., and Hanson, P.I. (2018). Triggered recruitment of ESCRT machinery promotes endolysosomal repair. Science 360, eaar5078.

Steele-Mortimer, O., Méresse, S., Gorvel, J.P., Toh, B.H., and Finlay, B.B. (1999). Biogenesis of Salmonella typhimurium-containing vacuoles in epithelial cells involves interactions with the early endocytic pathway. Cell. Microbiol. 1, 33–49.

Thurston, T.L.M., Wandel, M.P., Muhlinen, von, N., Foeglein, A., and Randow, F. (2012). Galectin 8 targets damaged vesicles for autophagy to defend cells against bacterial invasion. Nature 482, 414–418.

Wang, Y., Roche, O., Yan, M.S., Finak, G., Evans, A.J., Metcalf, J.L., Hast, B.E., Hanna, S.C., Wondergem, B., Furge, K.A., et al. (2009). Regulation of endocytosis via the oxygen-sensing pathway. Nat Med 15, 319–324.

Zhang, Z., Zhang, T., Wang, S., Gong, Z., Tang, C., Chen, J., and Ding, J. (2014). Molecular mechanism for Rabex-5 GEF activation by Rabaptin-5. Elife 3, e02687–e02687.

Zheng, Y.T., Shahnazari, S., Brech, A., Lamark, T., Johansen, T., and Brumell, J.H. (2009). The adaptor protein p62/SQSTM1 targets invading bacteria to the autophagy pathway. J. Immunol. 183, 5909–5916.

Zhu, G., Zhai, P., He, X., Wakeham, N., Rodgers, K., Li, G., Tang, J., and Zhang, X.C. (2004). Crystal structure of human GGA1 GAT domain complexed with the GAT-binding domain of Rabaptin5. EMBO J 23, 3909– 3917.

